# YAP1 oncogene is a context-specific driver for pancreatic ductal adenocarcinoma

**DOI:** 10.1101/270314

**Authors:** Bo Tu, Jun Yao, Sammy Ferri-Borgogno, Jun Zhao, Shujuan Chen, Qiuyun Wang, Liang Yan, Xin Zhou, Cihui Zhu, Seungmin Bang, Qing Chang, Christopher A. Bristow, Ya’an Kang, Hongwu Zheng, Huamin Wang, Jason B. Fleming, Michael Kim, Timothy P. Heffernan, Giulio F. Draetta, Duojia Pan, Anirban Maitra, Wantong Yao, Sonal Gupta, Haoqiang Ying

## Abstract

Transcriptomic profiling classifies pancreatic ductal adenocarcinoma (PDAC) into several molecular subtypes with distinctive histological and clinical characteristics. However, little is known about the molecular mechanisms that define each subtype and their correlation with clinical outcome. Mutant KRAS is the most prominent driver in PDAC, present in over 90% of tumors, but the dependence of tumors on oncogenic KRAS signaling varies between subtypes. In particular, squamous subtype are relatively independent of oncogenic KRAS signaling and typically display much more aggressive clinical behavior versus progenitor subtype. Here, we identified that YAP1 activation is enriched in the squamous subtype and associated with poor prognosis. Activation of YAP1 in progenitor subtype cancer cells profoundly enhanced malignant phenotypes and transformed progenitor subtype cells into squamous subtype. Conversely, depletion of YAP1 specifically suppressed tumorigenicity of squamous subtype PDAC cells. Mechanistically, we uncovered a significant positive correlation between WNT5A expression and the YAP1 activity in human PDAC, and demonstrated that WNT5A overexpression led to YAP1 activation and recapitulated YAP1-dependent but Kras-independent phenotype of tumor progression and maintenance. Thus, our study identifies YAP1 oncogene as a major driver of squamous subtype PDAC and uncovers the role of WNT5A in driving PDAC malignancy through activation of the YAP pathway.

## Introduction

Pancreatic ductal adenocarcinoma (PDAC) is projected to be the second leading cause of cancer-related mortality by 2030 (1), and displays as a highly heterogeneous disease with complex genetic and molecular diversity. Although the majority of PDACs share near-ubiquitous mutations of the *KRAS* oncogene and the frequent inactivation of *TP53*, *SMAD4* and *CDKN2A* tumor suppressors, additional somatic mutations occur at low individual prevalence, suggesting diverse non-genetic mechanisms underlying PDAC progression (2). Recent large-scale transcriptomic analyses classified human PDAC into several molecular subtypes with distinctive histological and clinical characteristics (3–6). However, the molecular subtypes are not consistently associated with any somatic mutations or other genetically altered pathways (6), suggesting that the biological phenotypes of these subsets are driven by subtype-specific molecular mechanisms other than genetic alterations. Besides the ADEX/exocrine and immunogenic subtypes, which are likely defined by signatures derived from non-neoplastic cells (4, 6), the molecular signatures of cancer cells largely fall into two categories: the squamous/quasimesenchymal/basal-like and the progenitor/classical subtypes. The squamous subtype reproducibly exhibits the worst prognosis compared with the other subtypes (3, 5, 6). Although the essential role of *KRAS* oncogene in tumor initiation and maintenance has been well appreciated (7, 8), it has been recently demonstrated that *KRAS* is dispensable for the survival of squamous subtype tumors (5, 9), suggesting that additional oncogenic drivers define and contribute to the malignancy of this subtype. Identifying the oncogenic pathways that drive the squamous subtype tumors will reveal subtype-specific vulnerabilities to treat these highly malignant tumors.

Yes-associated protein 1 (YAP1), is a transcriptional coactivator and plays critical roles in controlling normal tissue growth as well as tumor development (10). Its activity is kept in check by the upstream Hippo pathway, composed of the MST1/2-LATS1/2 kinases cascade, which phosphorylates YAP1 at multiple serine residues and sequesters YAP1 in cytoplasm for degradation (11). *In vivo* studies using genetically engineered mouse (GEM) models have shown that pancreas-specific *Yap1* depletion abolished PDAC development driven by oncogenic *Kra*s (12, 13), suggesting that YAP1 is essential for tumor initiation. However, the function of YAP1 in tumor maintenance in advanced human PDAC has not been firmly established. Notably, although we recently demonstrated that amplification of *Yap1* gene is capable of bypassing KRAS-dependency to maintain tumor growth in PDAC mouse model (14), the genetic alterations in the *YAP1* and core components of its upstream Hippo signaling pathway are very rare in human PDAC, pointing to a critical need to identify the nature of YAP expression and regulation as well as its association with clinic outcome in human PDAC.

In this study, we found that the YAP1 activation signature is highly enriched and preferentially required for the progression and maintenance of the squamous subtype PDAC. Gene expression profiling further uncovered a strong positive correlation of the non-canonical WNT pathway with the YAP1 activation signature; WNT5A, a prototypical non-canonical WNT ligand, is significantly upregulated in YAP1 activated tumors and also is required for YAP1 activation and tumorigenic activity in the squamous PDAC subtype. Moreover, we demonstrated that WNT5A enables the bypass of KRAS dependency to promote cell proliferation *in vitro* and drive tumor relapse *in vivo* in a YAP1-dependent manner. Together, our study delineated a critical role of the WNT5A-YAP1 axis in this deadliest form of human PDAC and identified context-specific vulnerabilities that may be exploited therapeutically.

## Results

### YAP1 plays a critical role in PDAC progression

We first evaluated the expression and role of YAP1 in human PDAC by using tissue microarray (TMA) analysis in a cohort of 92 human PDAC samples. As shown in Figure 1, 43 of 92 PDACs (47%) exhibited high YAP1 protein expression in tumor epithelium compared with the surrounding tissue; the median overall survival (OS) for the YAP1-low group was 38.3 months compared with 25.3 months for the YAP1-high group (*P* = 0.02) (Figure 1, A and B). Such association between elevated YAP1 protein and poor survival is similar to recent report (15) and was further validated in an independent cohort of 83 PDAC patients (*P* = 0.0475) (Figure 1C), suggesting that YAP1 may promote adverse biological outcomes in PDAC. We further characterized the *in vivo* function of YAP1 using GEM models. To faithfully recapitulate the PDAC initiation of human patients and investigated the requirement of YAP1 for PDAC when tumor initiation had started in the adult pancreas, we generated a tamoxifen-inducible YAP1 knockout mouse model of PDAC. Tamoxifen-induced acinar-specific activation of Cre recombinase in adult pancreas of the *Mist1-Cre^ERT2^*; *LSL-Kras^G12D/+^*; *LSL-Trp53^R172H/+^* (MKP) model led to rapid PDAC development accompanied by induction of nuclear YAP1 expression in tumor cells (Supplementary Figure 1A, *left*; YAP-WT). In contrast, acinar-specific deletion of *Yap1* in the MKPY (YAP-KO) model completely blocked tumor development (Supplementary Figure 1, A-C). Although all MKP mice succumbed to PDAC with a median survival of 103 days (n = 16), the *Yap1*-null MKPY mice remained entirely free of any overt pathological lesion (n = 13) (Supplementary Figure 1D). This could be attributed to the effect of YAP1 on cell proliferation and survival, as evidenced by the loss of both Ki67 and Survivin (BIRC5) staining in YAP-KO pancreas (Supplementary Figure 1, E and F). The protective effect of *Yap1* deletion on *Kras^G12D^* and *Trp53^R172H/+^* -induced PDAC development was further confirmed with an independent tamoxifen-inducible *Elastase-Cre^ERT2^*; *LSL-Kras^G12D/+^*; *LSL-Trp53^R172H/+^* model (Supplementary Figure 1G). These results suggested that YAP1 plays a critical role in PDAC progression.

**Figure 1.**
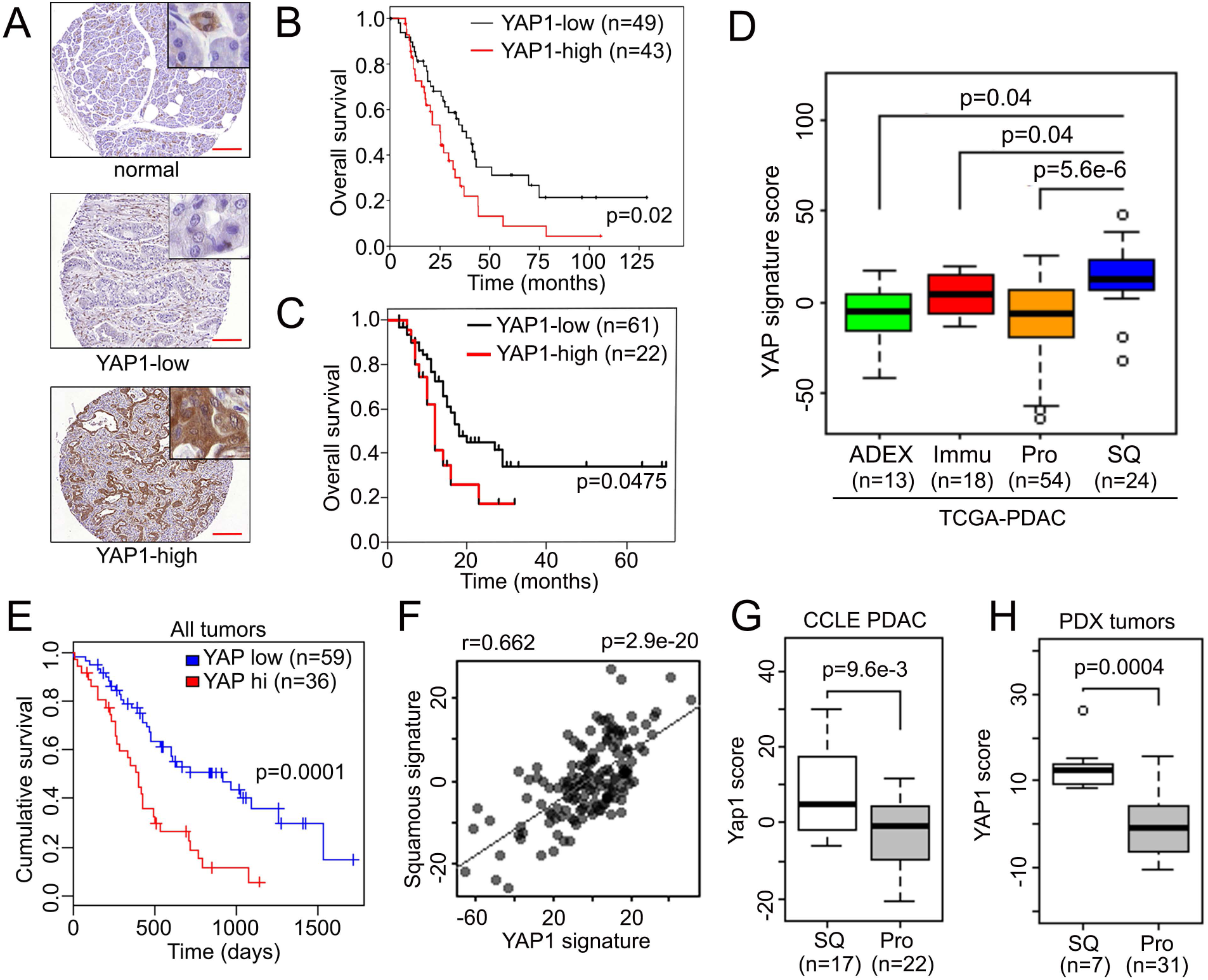
YAP1 is activated in the squamous subtype of human PDAC. **(A)** Representative images of YAP1 TMA showing tumor adjacent normal pancreatic tissue or tumor samples with low/high YAP1 level. Scale bar: 200 µm. **(B-C)** Kaplan-Meier curves for overall survival in PDAC patients from MD Anderson (MDA) **(B)** or Johns Hopkins University School of Medicine (JHU) (C) stratified by YAP1 expression. **(D)** YAP1 signature score among human PDAC subtypes in TCGA dataset. ADEX: aberrantly differentiated endocrine exocrine subtype; lmmu: immunogenic subtype; Pro: progenitor subtype; SQ: squamous subtype. **(E)** Kaplan-Meier curves for overall survival in all PDAC patients from TCGA dataset stratified by YAP1 activation signature score. **(F)** Correlation between squamous subtype signature and YAP1 signature in PDAC TCGA dataset. **(G)** YAP1 signature score in squamous (SQ) or progenitor (Pro) subtype human PDAC cell lines or PDXs **(H).** Error bars from all panels indicate ±SD. Pvalue for survival analysis was calculated with log rank test.

### YAP1 is activated in the squamous subtype of human PDAC

To further study the role of YAP1 in advanced human PDAC, we analyzed the expression profiles of the distinct molecular subtypes of human PDAC from the TCGA collection (4) and found that tumors of the squamous subtype exhibit elevated expression of genes that are known to be associated with YAP1 activation (16) (Figure 1D and Supplementary Table 1). Consistent with the TMA analysis, YAP1 pathway activation was significantly correlated with poor survival in PDAC patients (Figure 1E). Moreover, expression of the YAP1 activation signature was significantly correlated with that of the squamous subtype signature (Figure 1F), underscoring the tight association between YAP1 activation and squamous subtype tumors.

To further exclude the possibility that the YAP1 activation signature in squamous subtype PDAC is largely derived from tumor stroma, we analyzed the transcriptome data of human PDAC cell lines from the Cancer Cell Line Encyclopedia (CCLE) dataset and that of a collection of 47 PDAC PDX models, after the expression reads from murine host were omitted. Consistent with the notion that the molecular signatures of ADEX or immunogenic subtypes are likely derived from non-tumor cells (4), we failed to identify these signatures in both human PDAC cell lines and PDX (Supplementary Figure 2, A and B and data not shown). Not surprisingly, the molecular signatures of human PDAC cell lines or PDXs clustered primarily under either the progenitor or squamous subtype (Supplementary Figure 2, A and B). In accordance with analysis of TCGA data, the YAP1 activation signature was consistently elevated in the squamous subtype cells (Figure 1, G and H), underscoring that YAP1 is preferentially activated in the squamous PDAC subtype.

**Figure 2.**
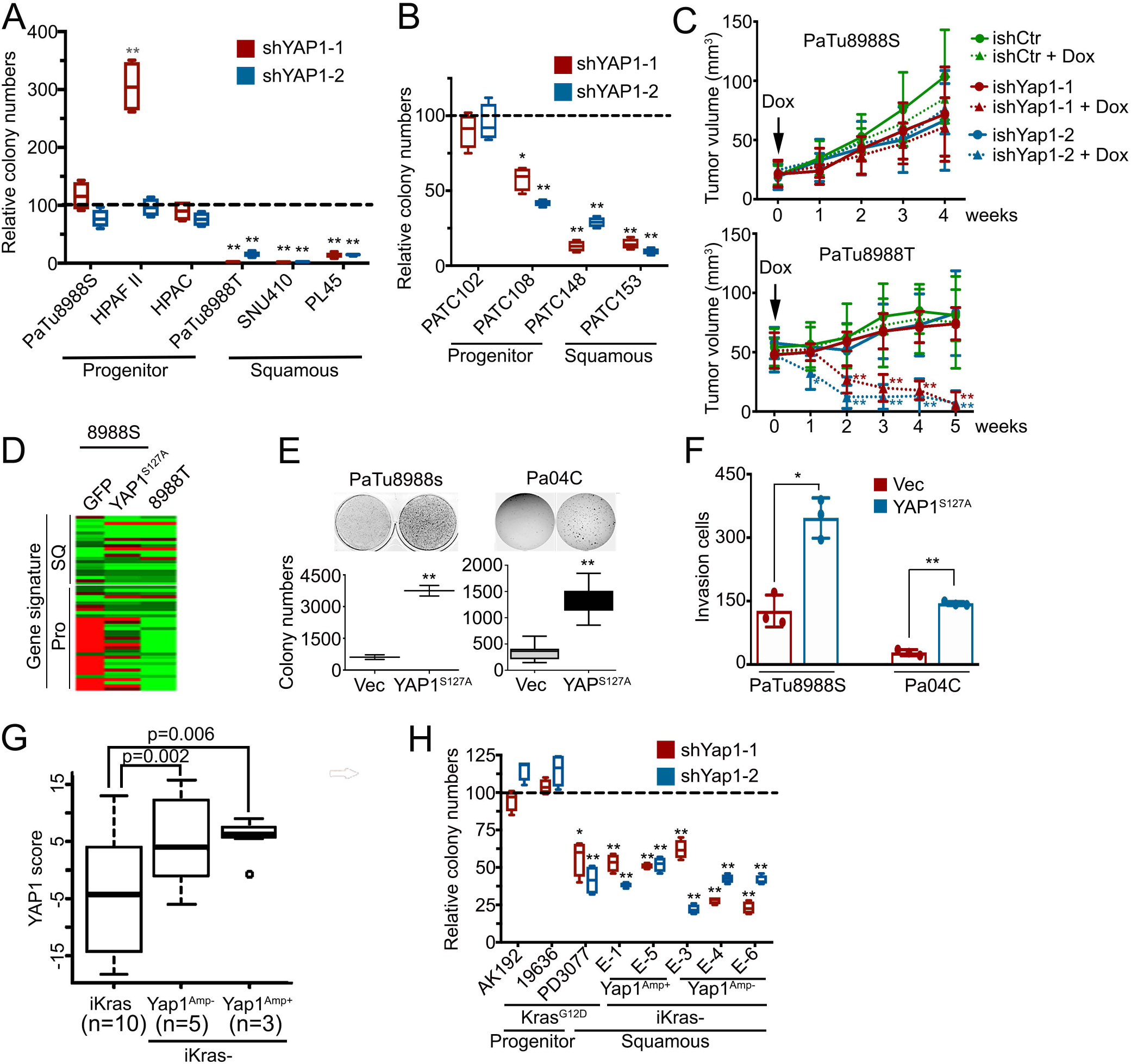
YAP1 is essential for the maintenance of squamous subtype of PDACs. **(A-B)** Colony formation assay in human PDAC eell lines **(A)** or POX cells **(B)** infected with YAP1 shRNAs or nontargeting shRNA. Relative colony numbers upon normalization to the nontargeting shRNA group is shown (n=3). **(C)** PaTu8988S and PaTu8988T cells engineered with inducible shRNA targeting YAP1 or non-targeting shRNA were subcutaneously injected into nude mice. Animals were treated with doxycycline once tumor size reached 20-50 mm^3^. Tumor volumes were measured on the indicated dates post-injection (n=5). **(D)** Heatmap shows the expression level of squamous (SQ) and progenitor (Pro) subtype signature genes measured with NanoString in PaTu8988S cells expressing GFP or YAP1^S127A^ and PaTu8988T cells. **(E-F)** Ectopic expression of YAP1^S127A^ in PaTu8988S and Pa04C cells promotes anchorage-independent growth in soft agar (quantification from triplicates shown in bottom panel) **(E)** and cell invasion in a Boyden chamber assay (Quantification of triplicates is shown) **(F). (G)** YAP1 signature score among mouse PDAC tumors with indicated genotypes. **(H)** Colony formation assay in mouse PDAC cells infected with YAP1 shRNAs or nontargeting shRNA. Relative colony numbers upon normalization to the nontar­ geting shRNA group is shown (n=3). Error bars from all panels indicate ±SD. *: *P* < 0.05; **: *P* < 0.01.

### YAP1 is essential for the maintenance of the squamous subtype PDAC

To further investigate the requirement of YAP1 in the squamous subtype of PDAC, we conducted loss-of-function studies with shRNA in a panel of human PDAC cell lines and early-passage primary cell lines derived from human PDX tumors. Knockdown of YAP1 strongly suppressed the colony-formation capacity of PDAC cell lines (PaTu8988T, SNU410, and PL45) and PDX cell lines (PATC148 and PATC153) belonging to the squamous subtype with a strong YAP1 activation signature (Figure 2, A and B and Supplementary Figure 3, A and B). In contrast, the progenitor subtype cell lines, including PaTu8988S, HPAF-II, HPAC PATC102, and PATC108 cells, were relatively insensitive to YAP1 depletion (Figure 2, A and B and Supplementary Figure 3, A and B). Moreover, inducible knockdown of YAP1 in established tumors resulted in inhibition of proliferation, induction of apoptosis and eventual regression of PaTu8988T tumors, whereas the growth of PaTu8988S tumors was not affected (Figure 2C and Supplementary Figure 3, C-E), indicating the role of YAP1 for tumor maintenance in the squamous PDAC subtype.

**Figure 3.**
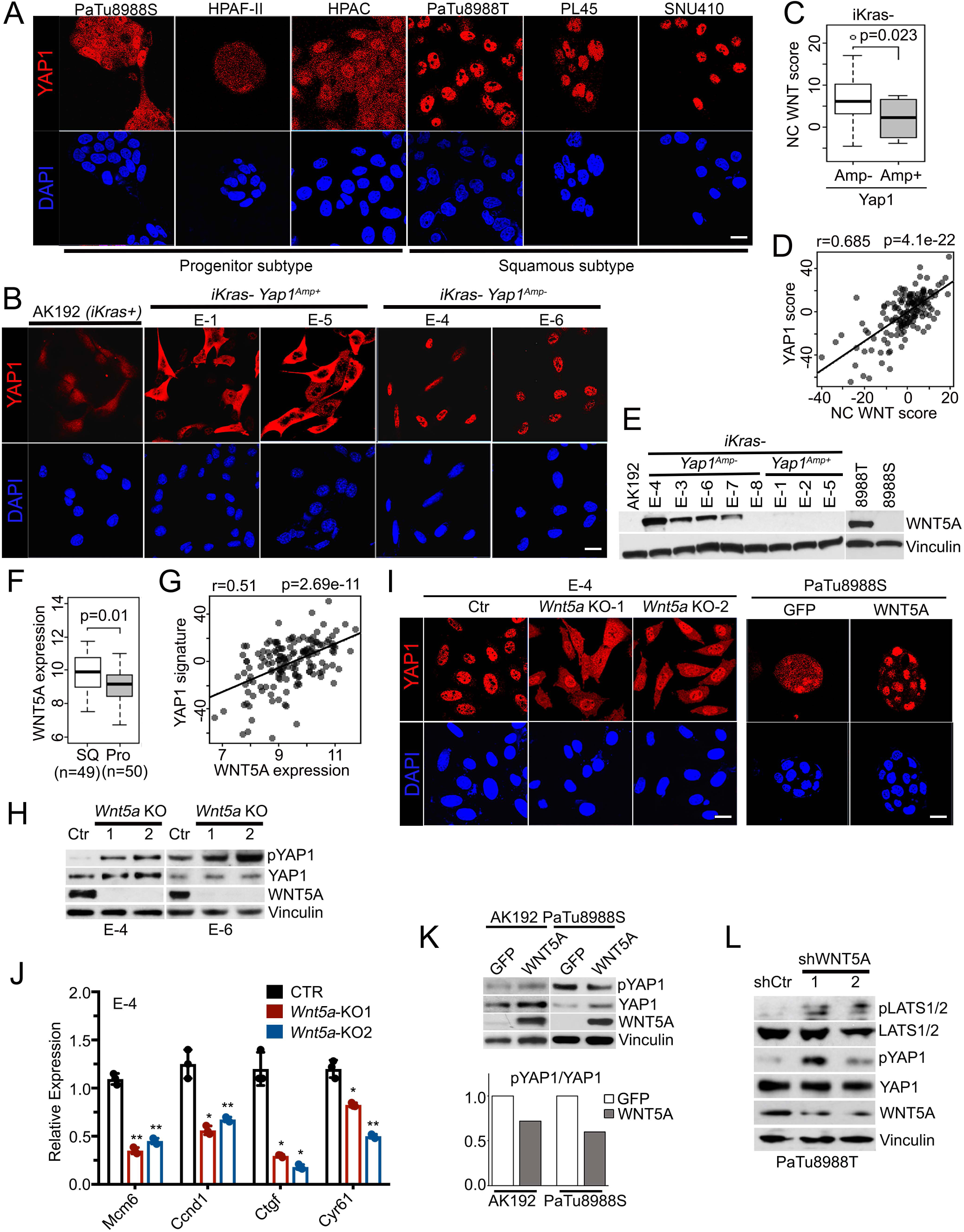
YAP1 activation in PDAC is mediated by WNT5A overexpression. **(A-B)** Human pancreatic cancer cell lines of progenitor or squamous subtype **(A)** or mouse PDAC cells of indicated genotypes **(B)** were subjected to immunofluorescence staining with anti-YAP1 (red) and DAPI (blue). **(C)** Noncanonical (NC) WNT pathway enrichment score in iKras-mouse PDAC tumors without (Amp-) or with (Amp+) *Yap1* amplification. **(D)** Correlation between NC WNT signature and YAP1 signature in PDAC TCGA dataset. **(E)** Western blots for WNT5A in mouse PDAC cells of indicated genotypes and human PDAC cell lines. **(F)** WNT5A expression in squamous (SQ) or progenitor (Pro) subtype human PDACs in TCGA dataset. **(G)** Correlation between WNT5A expression and YAP1 signature in PDAC TCGA dataset. **(H)** Western blot analysis for WNT5A, YAP1, and phospho-YAP1 (S127) in two independent *iKras-Yap1^Amp-^* cells with CRISPR-mediated Wnt5a deletion. Two independent *Wnt5a* knockout (KO) clones were included, and a clone without *Wnt5a* deletion was used as control (Ctr). **(I)** E-4 *(iKras-Yap1^Amp-^)-Wnt5a* KO cells and PaTu8988S cells expressing GFP or WNT5A were subjected to immunofluorescence staining with anti-YAP1 (red) and DAPI (blue). **(J)** Relative mRNA levels of YAP1 downstream targets in E-4 *(iKras-Yap1^Amp-^)-Wnt5a* KO cells. **(K)** Western blot analysis for WNT5A, YAP1, and phospho-YAP1 (S127) in mouse iKras PDAC cells (AK192) or human PaTu8988S cells expressing GFP or WNT5A (top). The quantification of phosphor-YAP1/total YAP1 signals is shown in the bottom panel. **(L)** Western blot analysis for WNT5A, LATS1/2, phospho-LATS1/2, YAP1, and phospho-YAP1 (S127) in PaTu8988T cells infected with WNT5A shRNAs

We further investigated whether YAP1 is able to endow progenitor subtype cells with the squamous phenotype. Expressing constitutive active YAP1^S127A^ mutant, which is resistant to cytoplasmic retention and degradation (11), in PaTu8988S cells, a progenitor subtype human PDAC cell line, induced a transcriptional signature that was highly similar to that of PaTu8988T cells, which were derived from the same patient as PaTu8988S cells (17) but of the squamous subtype (Figure 2D and Supplementary Table 2). Of importance, ectopic expression of YAP1^S127A^ substantially enhanced the anchorage-independent growth, migration, and invasion capacity of PaTu8988S cells and an early passage patient-derived cell line Pa04C (Figure 2, E and F and Supplementary Figure 3, F and G), suggesting active YAP1 may promote the tumorigenicity and metastatic spread of PDAC cells. Indeed, YAP1^S127A^ expression diminished necrotic regions within primary tumor core and enhanced the distal metastasis of Pa04C cells in an orthotopic xenograft model (Supplementary Figure 3, H-J). This was accompanied with induction of canonical YAP1 target genes, such as *CYR61*, *CTGF,* and *AXL*, in both cultured cells and xenograft tumors (Supplementary Figure 3, K and L). Gene expression microarray and subsequent Gene Set Enrichment Analysis (GSEA) in these cells confirmed an established YAP1 signature (Supplementary Figure 3M), enrichment in pathways associated with tumor development and metastasis, and the underlying cellular processes responsible, such as cell proliferation, cell cycle progression, migration, motility, and epithelial-to-mesenchymal transition (EMT) (Supplementary Tables 3 and 4). Of interest, cellular processes including cell cycle progression and signaling pathways significantly activated in Pa04C-YAP1^S127A^, such as MYC, IL6-STAT3, TGFβ, RhoA, E2F, and TNF, along with the Hippo signaling pathway, were part of all four gene signatures (GP2-5) associated with the squamous subtype of PDAC (3) (Supplementary Tables 5 and 6), underscoring the role of YAP1 activation in this most aggressive subtype.

We and others have previously shown that YAP1 activation enables the bypass of oncogene addiction in multiple cancer types, including PDAC (14, 18–20). Indeed, compared with PaTu8988S cells, PaTu8988T cells were more resistant to KRAS knockdown with shRNA (Supplementary Figure 4, A-C). Expression of YAP1^S127A^ partially rescued the growth of PaTu8988S cells upon KRAS depletion (Supplementary Figure 4, B and C), indicating that YAP1 activation could enable the bypass of KRAS dependence in human PDAC cells. Accordingly, pathway analysis of human PDAC expression profiles in the TCGA dataset indicated that gene signatures induced upon KRAS knockdown (21) or suppressed by oncogenic KRAS expression (22) were significantly upregulated in squamous subtype tumors (Supplementary Figure 4D), suggesting relatively low KRAS activity in these tumors. By using an inducible *Kras^G12D^*-driven PDAC GEM model, we recently obtained a collection of spontaneous relapse tumors following *Kras^G12D^* extinction in advanced PDAC (iKras– tumors) (14). These tumors neither expressed oncogenic *Kras* nor exhibited strong activation of KRAS surrogates and are thus deemed KRAS-independent. Of interest, the molecular signature of the iKras– tumors (Supplementary Table 7), which is composed of genes that are highly expressed in iKras-tumors compared with those in iKras+ ones (14), was also significantly enriched in the squamous subtype of human PDACs (Supplementary Figure 4, E and F), further supporting the notion that squamous subtype tumors are relatively KRAS-independent. Consistent with YAP1 activation in the human squamous subtype PDAC, the YAP1 signature was significantly enriched in mouse iKras– tumors (Figure 2G). Although YAP1 knockdown exhibited minimal effect on progenitor subtype *Kras*-driven mouse tumor lines (AK192 and 19636), YAP1 depletion significantly suppressed the proliferation and colony formation capability of squamous subtype cells (PD3077 (23)) and iKras– tumor cells (Figure 2H and Supplementary Figure 4G), underscoring YAP1 dependency in the squamous subtype of tumors.

**Figure 4.**
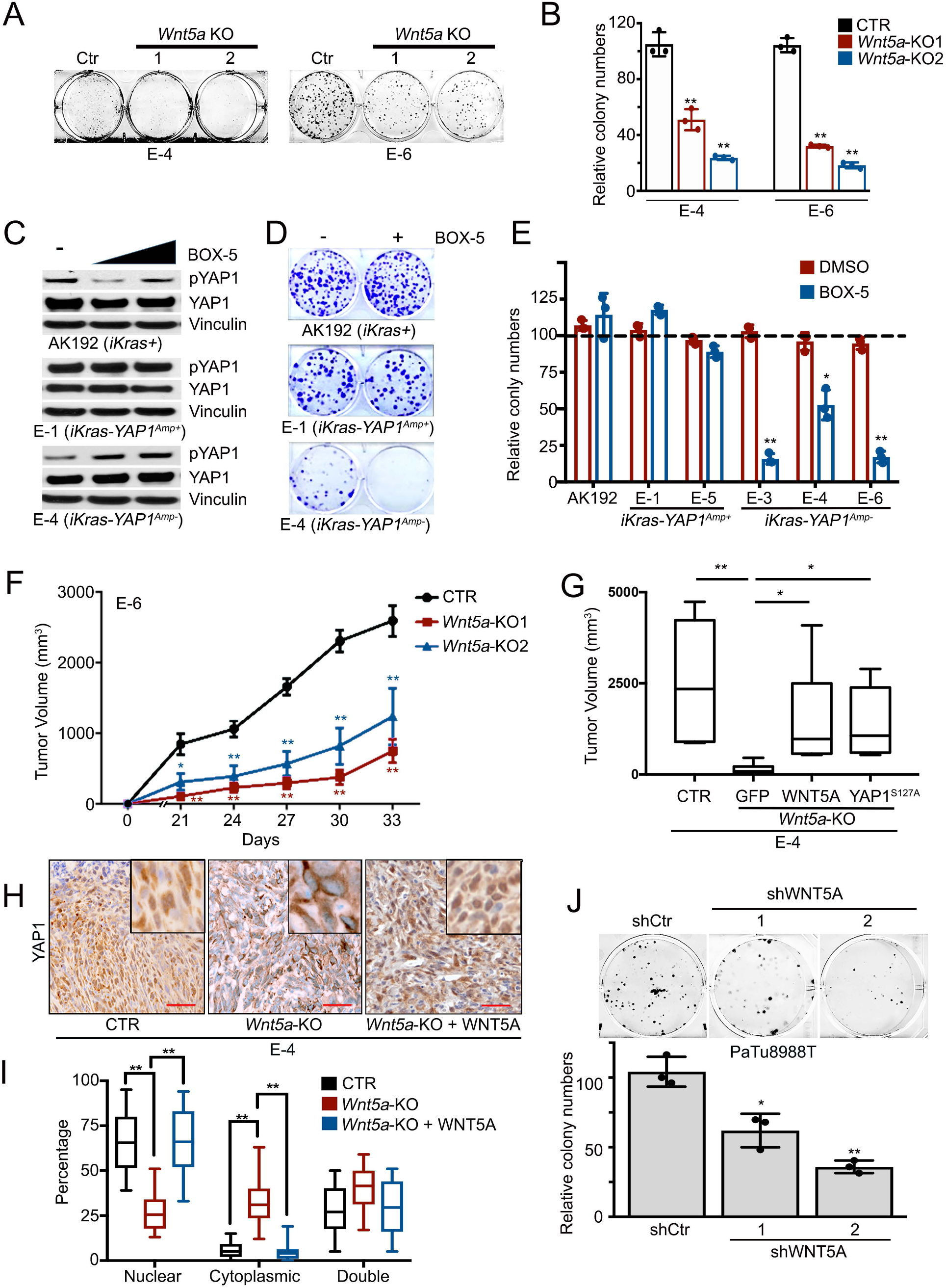
WNT5A overexpression is required for tumorigenic activity. **(A)** Representative images of the colony formation assay for *iKras-Yap1^Amp-^-Wnt5a* KO cells. Clones without *Wnt5a* deletion were used as a control (Ctr). Quantification from triplicates is shown in **(B)** and is pre­ sented as relative colony numbers upon normalization to the shCtr group. **(C)** Western blot analysis for YAP1 and phospho-YAP1 (S127) in mouse PDAC cells treated with DMSO, 50 or 100 µM BOX-5. **(D)** Representative images of the colony formation assay in mouse PDAC cells of indicated genotypes treated with vehicle (DMSO) or BOX-5 (100 µM). Quantification from triplicates is shown in **(E)** and is presented as relative colony numbers upon normalization to DMSO group. **(F)** Two independent clones of E-6 *(iKras-Yap1^Amp-^)-Wnt5a* KO cells and control cells without *Wnt5a* deletion (CTR) were subcutaneously injected into nude mice. Tumor volumes were measured on the indicated dates after injection. Results are presented as the means± SD (n = 5). **(G)** E-4 *(iKras-Yap1^Amp-^)-Wnt5a* KO cells were infected with GFP, WNT5A, or YAP1S127A and were subcutaneously injected into nude mice. Cells without Wnt5a deletion were used as a control (CTR). Tumor volumes were measured 30 days after injection. Results are presented as the means ± SD (n = 5). **(H)** Subcutaneous xenograft tumors from **(G)** were stained for YAP1. Scale bar: 100 µm. Percentage of cells with nuclear/cytoplasmic/double staining of YAP1 is shown in (I). Error bars represent SD (n = 10 fields, 250 cells/field). **(J)** Representative images of the colony formation assay for PaTu8988T cells infected with WNT5A shRNAs or nontargeting shRNA (shCtr) (top). Quantification from triplicates is shown

### WNT5A overexpression contributes to YAP1 activation in PDAC

In an effort to identify mechanism for YAP1 activation, we first performed genomic analysis of primary human tumors from the TCGA dataset, showing that no frequent copy number changes or mutations of *YAP1* locus were revealed (Supplementary Figure 5A). Similarly, although *Yap1* is amplified in a subset of mouse iKras– tumors (14), most iKras– tumors harbor no genomic alteration of *Yap1* despite enrichment of the YAP1 signature (Figure 2G and Supplementary Figure 5B). In addition, there is no obvious difference in YAP1 transcription level between squamous and progenitor subtype of PDAC cell lines (Supplementary Figure 5C), indicating post-transcriptional mechanisms for YAP1 activation in PDAC. Indeed, YAP1 activation as manifested by its nuclear translocation (11) was evident in YAP1-dependent human PDAC cells, including PaTu8988T, SNU410, and PL45 cells, and iKras– mouse tumors or the derived cell lines without *Yap1* amplification (*iKras*– *Yap1^Amp-^*) (Figure 3, A and B and Supplementary Figure 5D). Conversely, YAP1 was localized in both the cytoplasm and nuclei in PDAC cells with weak YAP1 signature, such as PaTu8988S, HPAF-II, and HPAC cells, as well as progenitor subtype *Kras^G12D^*-driven mouse tumors and iKras– tumors with *Yap1* amplification (*iKras*– *Yap1^Amp+^*) (Figure 3, A and B). Accordingly, YAP1-activated human and mouse PDAC cells exhibit reduced phosphorylation at S127 (Supplementary Figure 5, E and F), suggesting that YAP1 activation is largely mediated by upstream Hippo pathway at the posttranslational level.

**Figure 5.**
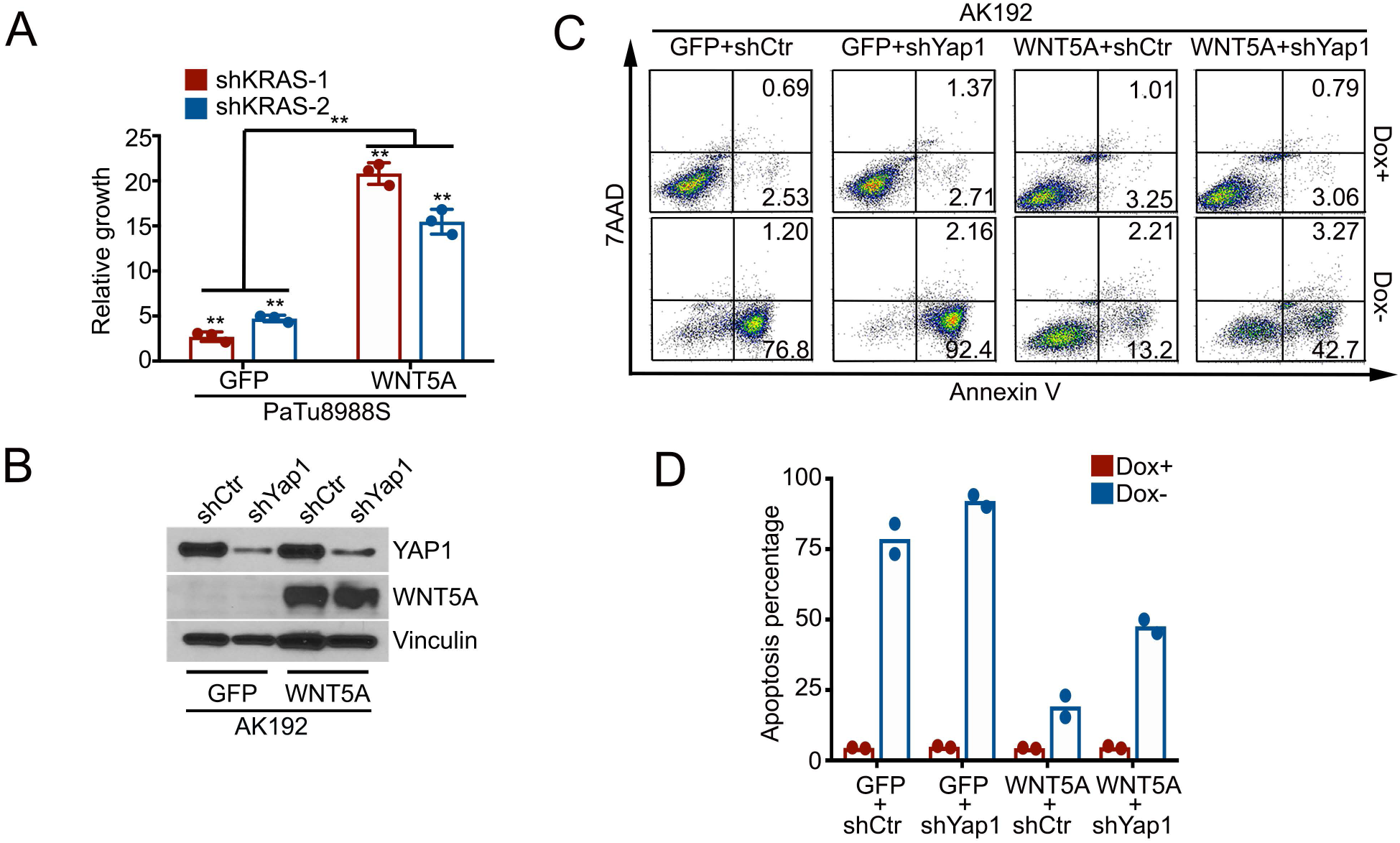
WNT5A overexpression leads to the bypass of KRAS dependency. **(A)** Cell growth assay for PaTu8988S-GFP or -WNT5A cells infected with KRAS shRNAs or nontargeting control shRNA. Quantification from triplicates is presented as relative cell growth upon normalization to control group. Error bars indicate ±SD of triplicates, **: *P* < 0.01. **(B)** Western blot analysis for WNT5A and YAP1 in AK192 *(iKras+)* cells expressing GFP or WNT5A upon knockdown of YAP1 with shRNA. **(C)** Mouse AK192-GFP or AK192-WNT5A cells infected with nontargeting shRNA (shCtr) or YAP1 shRNA (shYap1) were grown as 3D tumor spheres in the presence or absence of doxycycline for 4 days. Cellular apoptosis was measured with annexin V staining. Representative images of two independent experiments show the FACS analysis of annexin V and 7MD staining. Numbers represent the percentage of early apoptosis (Annexin v^+^ 7AAD^-^) and late apoptosis (annexin v^+^ 7AA^+^) populations. **(D)** Quantification of total apoptotic cell from **(C)** (two independent experiments). Doxycycline withdrawal leads to dramatic apoptosis of AK192-GFP tumor sphere. Such apoptosis induced by doxycycline withdrawal was significantly inhibited in WNT5A-expressing cells, which was partially reversed upon YAP1 knockdown.

We further performed GSEA analysis to compare the differentially regulated pathways in *iKras*– *Yap1^Amp-^* vs *iKras*– *Yap1^Amp+^* cells. Of interest, one of the pathways significantly elevated in *iKras*– *Yap1^Amp-^* cells was the noncanonical WNT pathway (24) (Figure 3C and Supplementary Table 8), which was recently showed to suppress Hippo signaling and activate YAP1 in adipocytes (25). Of importance, a significant correlation between the noncanonical WNT signature and YAP1 activation signature was observed in the TCGA dataset (Figure 3D), suggesting control of YAP1 activation by the noncanonical WNT pathway in PDAC. A survey of WNT ligands identified *Wnt5a*, the prototypic noncanonical WNT ligand (26), to be exclusively overexpressed in the *iKras*– *Yap1^Amp-^* tumor cells at both mRNA and protein levels (Figure 3E and Supplementary Figure 5G). Moreover, *WNT5A* expression was also significantly elevated in the squamous subtype compared with its expression in progenitor subtype tumors and was significantly correlated with the YAP1 signature in the TCGA dataset (Figure 3, F and G). Deletion of *Wnt5a* with CRISPR in two independent *iKras*– *Yap1^Amp-^* tumor cell lines led to the induction of YAP1 phosphorylation at S127, the increase in cytoplasmic retention of YAP1 protein, as well as the downregulation of YAP1 downstream target genes (Figure 3, H–J). In addition, ectopic expression of *Wnt5a* in *Kras^G12D^*-driven tumor cells resulted in a decrease in YAP1 phosphorylation (Figure 3K), further supporting the notion that WNT5A expression drives YAP1 activation in mouse PDAC cells. We further validated these findings in human PDAC cell lines, in which WNT5A expression was found to be highly elevated in the YAP1-dependent PaTu8988T cells in contrast to progenitor subtype PaTu8988S cells (Figure 3E). Ectopic expression of WNT5A in PaTu8988S cells reduced YAP1 phosphorylation and enhanced YAP1 nuclear localization (Figure 3, K and I). In contrast, depletion of WNT5A in PaTu8988T cells with shRNA resulted in elevated YAP1 phosphorylation with concordant induction of LATS1/2 phosphorylation (Figure 3L), supporting the notion that WNT5A activates YAP1 through the suppression of Hippo signaling. Taken together, our data suggest that WNT5A overexpression can lead to YAP1 activation in PDAC cells.

### WNT5A overexpression enables tumor maintenance and bypass of KRAS dependence

At the functional level, *Wnt5a* deletion in *iKras*– *Yap1^Amp-^* tumor cells with CRISPR significantly inhibited colony formation (Figure 4, A and B). In agreement with the genetic ablation, treatment with WNT5A antagonist BOX-5 specifically induced YAP1 phosphorylation and abolished the colony-formation ability of *iKras*– *Yap1^Amp-^* tumor cells, but not the *Yap1^Amp+^* cells (Figure 4, C– E), indicating the requirement of WNT5A for YAP1 activation and the induced tumorigenic activity. Indeed, *Wnt5a* deletion also significantly inhibited xenograft tumor growth *in vivo*, which was rescued by reconstituted WNT5A expression (Figure 4, F and G). Of importance, in contrast to the predominant nuclear staining of YAP1 in the parental *iKras*– *Yap1^Amp-^* cells, *Wnt5a* knockout tumors exhibited a significant amount of cytoplasmic YAP1, whereas WNT5A reconstitution restored YAP1 nuclear accumulation without affecting total YAP1 expression level (Figure 4, H and I and Supplementary Figure 6A), suggesting that the diminished tumor growth was due to decreased YAP1 activity. Consistent with the role of YAP1 in driving squamous subtype, expression of squamous subtype genes (Cav1 and Pappa) was suppressed in *Wnt5a* KO tumors whereas the expression of progenitor subtype genes (Tff1 and Muc13) was induced, which was partially reversed upon WNT5A reconstitution (Supplementary Figure 6B). In addition, expression of constitutively active YAP1^S127A^ largely rescued the inhibitory effect of *Wnt5a* deletion on tumor growth (Figure 4G), thus suggesting that WNT5A overexpression in mouse PDAC cells promotes tumor growth by activating YAP1. In agreement with this notion, depletion of WNT5A in human PDAC cell line PaTu8988T significantly inhibited cell colony formation ability (Figure 4J).

Since YAP1 activation can maintain tumor growth upon genetic extinction of KRAS oncogene in PDAC (14, 20), we next investigated whether WNT5A overexpression can also serve to bypass KRAS dependency. Indeed, ectopic expression of WNT5A in *Kras^G12D^*-driven iKras tumor cells and KRAS-dependent PaTu8988S cells partially restored the colony formation upon KRAS depletion (Figure 5A and Supplementary Figure 6, C and D). In addition, forced WNT5A expression in *Kras^G12D^*-driven iKras tumor spheres was able to maintain cell viability upon extinction of *Kras^G12D^* by doxycycline withdrawal, whereas most control tumor cells expressing GFP underwent apoptosis (Figure 5, B–D). Of importance, the survival effect of WNT5A upon *Kras^G12D^* extinction was largely abolished upon YAP1 knockdown (Figure 5, B–D), indicating that YAP1 is required for WNT5A-induced bypass of KRAS dependence. Indeed, similar to the effect of YAP1^S127A^, ectopic WNT5A expression in iKras tumor cells showed KRAS-independent tumor growth when injected orthotopically into nude mice, whereas GFP-expressing iKras tumor cells failed to maintain tumor growth in the absence of doxycycline (Figure 6, A–C). Of importance, WNT5A-induced KRAS-independent tumor growth was abolished in *Yap1*-deleted cells (Supplementary Figure 6, E and F), underlying the requirement of YAP1 for WNT5A-mediated bypass of KRAS dependence. Notably, all WNT5A-driven tumors showed lower MAPK activity and strong nuclear YAP1 accumulation compared with *Kras^G12D^*-driven tumors (Figure 6D). Together, these results indicate that WNT5A overexpression can activate YAP1 and substitute for oncogenic Kras-driven tumor maintenance.

**Figure 6.**
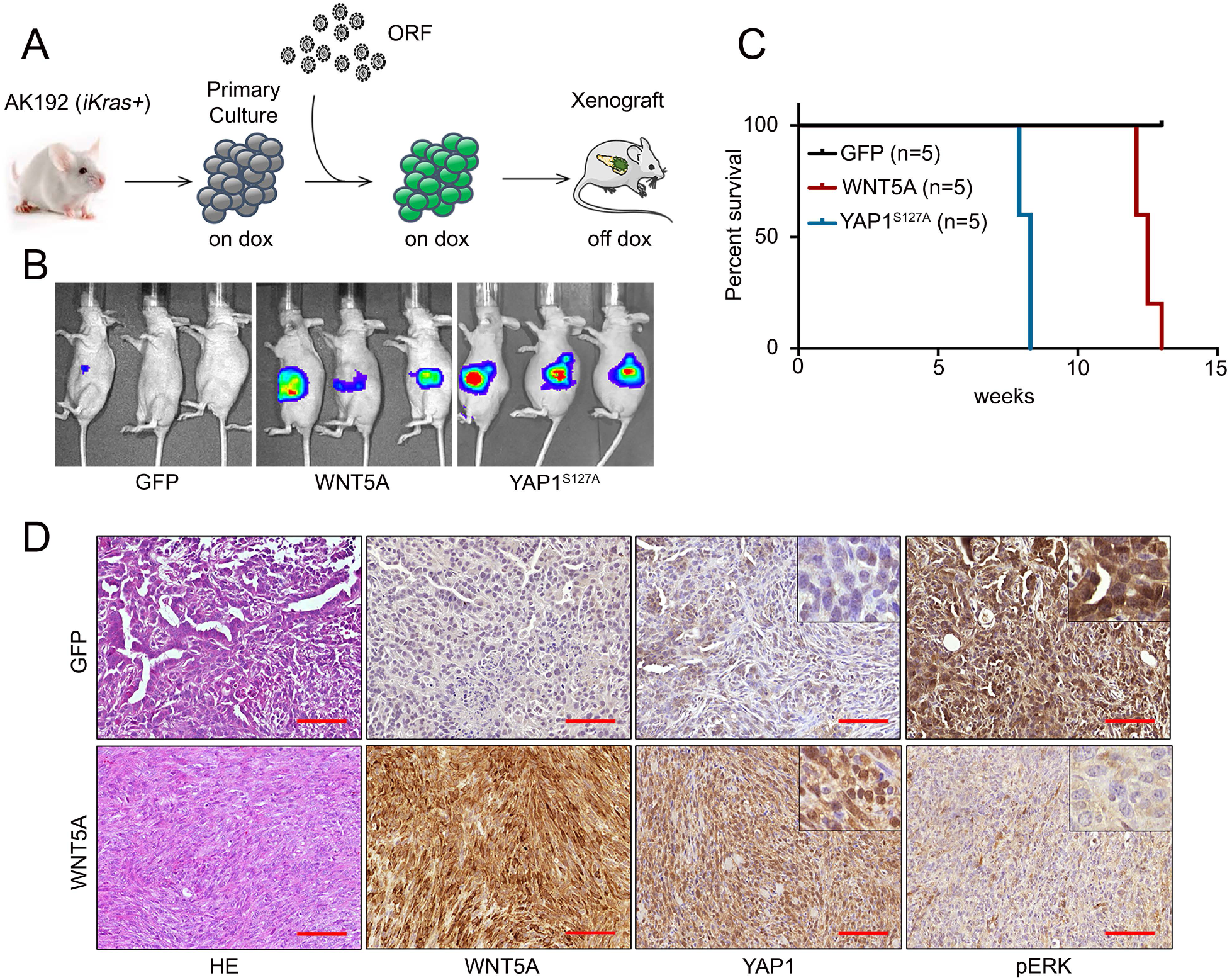
WNT5A overexpression drives KRAS-independent tumor growth. **(A)** Schematic workflow for the in vivo KRAS bypass experiment. **(B)** AK192 cells expressing luciferase were infected with lentivirus expressing GFP, WNT5A, or YAP1S127A and orthotopically injected into nude mice pancreas in the presence of doxycycline. Animals were withdrawn from doxycycline 4 days later, and tumor growth was visualized by bioluminescent imaging at 8 weeks. **(C)** Kaplan-Meier overall survival analysis for nude mice (n = 5/group) orthotopically transplanted with the cells described in **(B). (D)** Orthotopic xenograft tumors generated with AK192-GFP cells (on doxycycline) or AK192-WNT5A cells (off doxycycline) were stained for WNT5A, YAP1, and phospho-ERK. Scale bar: 100 µm.

### WNT5A-YAP1 axis functions in primary human PDAC

We further validated the WNT5A-YAP1 axis in primary human PDAC, showing that *WNT5A* expression was elevated in squamous subtype PDX tumors (Figure 7A). Accordingly, WNT5A protein was highly expressed, whereas YAP1 phosphorylation was relatively low in squamous subtype PATC148 and PATC153 cells, while two cell lines derived from progenitor subtype tumors, PATC102 and PATC108, exhibited elevated YAP1 phosphorylation along with absence of WNT5A expression (Figure 7B). This is in accordance with the elevated expression of YAP1 target gene CYR61 in squamous subtype PDXs (Figure 7B).

**Figure 7.**
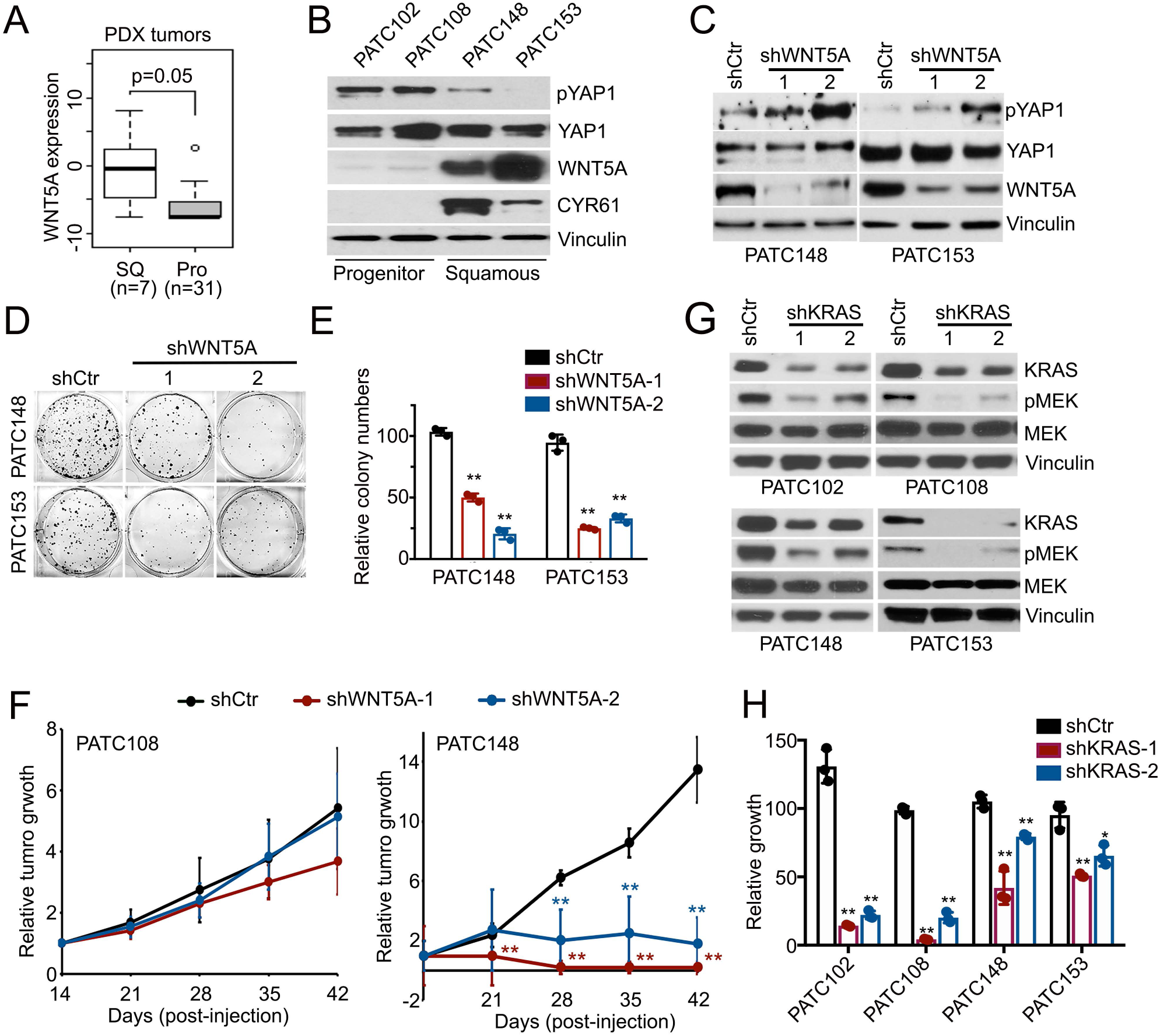
WNT5A-YAP1 axis is active in primary human PDAC and required for tumor maintenance. **(A)** WNT5A expression in squamous (SQ) or progenitor (Pro) subtype PDXs. **(B)** Western blot analysis for WNT5A, YAP1, phospho-YAP1 (S127), and CYR61 in POX cell lines of squamous or progenitor subtype. **(C)** Western blot analysis for WNT5A, YAP1, and phospho-YAP1 (S127) in squamous subtype POX cell lines infected with WNT5A shRNAs or nontargeting shRNA (shCtr). **(D)** Representative images of the colony formation assay for squamous subtype POX cell lines infected with WNT5A shRNAs or nontargeting shRNA (shCtr). Quantification from triplicates is shown in **(E). (F)** PATC108 and PATC148 cells infected with WNT5A shRNAs or nontargeting shRNA (shCtr) were subcutaneously injected into nude mice. Tumor volumes were measured on the indicated dates after in­ jection, and relative tumor growth normalized to day 14 was presented. Results are presented as the means± SD (n = 5). **(G)** Western blot analysis for KRAS, phosphor-MEK1/2, and MEK1/2 in POX cell lines infected with KRAS shRNAs or nontargeting shRNA (shCtr). **(H)** Cell growth assay for POX cell lines infected with KRAS shRNAs or nontargeting shRNA (shCtr). Quantification from triplicates is presented as relative cell growth upon nor­ malization to the shCtr group. Error bars from all panels indicate ±SD. *: *P* < 0.05; **: *P* < 0.01.

ShRNA-mediated depletion of WNT5A in PDX-derived PATC148 and PATC153 cells caused an increase in YAP1 phosphorylation, suppression of colony-formation ability, and diminished tumor growth *in vivo* (Figure 7, C–F), supporting the role of WNT5A for tumorigenic activity. In contrast, WNT5A shRNA had minimal effect on *in vivo* tumor growth of PATC108 cells (Figure 7F). Consistent with the role of WNT5A in bypass of KRAS dependence, knockdown of KRAS elicited less inhibition on the growth of high WNT5A expressing PATC148 (*KRAS^G12D^*) cells compared with low WNT5A expressing PATC102 (*KRAS^G12D^*) and PATC108 (*KRAS^G12D^*) cells, with *KRAS WT* PATC153 cells being resistant to KRAS knockdown (Figure 7, G and H). Together, our data indicate that WNT5A overexpression in squamous subtype PDACs contributes to YAP1 activation and tumor growth.

## Discussion

In this study, we found that deletion of *Yap1* in adult pancreas completely blocked KRAS-induced PDAC development (Supplementary Figure 1, B–D). Immunohistochemical staining on these tumor tissues showed marked decrease in the cell proliferation index, as measured by Ki-67 staining (Supplementary Figure 1F) and low Survivin (BIRC5) expression (Supplementary Figure 1E). Since Survivin expression overlapped with YAP1 expression in *Yap1*-WT PDAC but was completely lost in *Yap1*-KO pancreas, it is likely regulated directly by YAP1 at the transcription level, as shown in esophageal squamous cell carcinoma (27). Survivin is a known anti-apoptotic protein, which is expressed only in tumor cells (28) and primarily during the G2-mitotic phase of the cell cycle (29). The mostly nuclear expression of Survivin that is observed in YAP-WT tumor sections (Supplementary Figure 1E) supports its predominant role in regulating the cell cycle. Accordingly, higher Survivin expression was also in agreement with predominant gene signatures associated with cell cycle progression in Pa04C-YAP1^S127A^ cells (Supplementary Table 4) and a higher percentage of Pa04C-YAP1^S127A^ cells in the G2/M phase by cell cycle analysis (data not shown).

We provided evidence that YAP1 is highly activated in squamous subtype PDACs and required for their tumorigenic function. Despite the emerging role of *YAP1* as a major oncogene in multiple cancer types, genetic alterations of the *YAP1* gene or its upstream Hippo pathway are relatively uncommon (30, 31). YAP1 amplification or mutations in *NF2*, an upstream negative regulator of YAP1 activity, has been reported in about 1% of human PDAC (3, 32). Therefore, the activation of YAP1 in advanced human PDAC is likely due to nongenetic factors regulating inhibitory upstream Hippo kinases. Our data showing constitutive nuclear localization of YAP1 protein in YAP1-dependent tumor cells indicates that suppression of Hippo signaling is the major mechanism for YAP1 activation in PDAC. It was shown that TP63 (ΔNp63) drives the squamous subtype of PDAC (33). Of interest, TP63 has been shown to activate YAP1 in head and neck squamous cell carcinoma through the suppression of Hippo signaling (34). Whether TP63 also functions through YAP1 activation in the squamous subtype of PDAC remains to be investigated. More recently, GLI2 transcriptional factor was demonstrated to drive the switch between progenitor and squamous subtypes in PDAC (35). Importantly, GLI2 was identified as a direct downstream target of YAP1 in medulloblastoma (36). It will be interesting to further investigate whether YAP1 also functions upstream of GLI2 in squamous subtype PDAC.

Here we provide evidence that WNT5A overexpression leads to YAP1 activation and bypass of KRAS dependency in KRAS-independent mouse PDAC cells and a subset of human squamous subtype of PDACs. WNT5A is a prototypic noncanonical WNT ligand (26) and has been implicated in the pathogenesis of PDAC (37, 38). It was recently showed that the WNT5A-mediated noncanonical WNT pathway suppresses Hippo signaling and activates YAP1 through G protein-dependent activation of Rho GTPases (25). WNT5A can also engage multiple additional downstream signaling pathways, including SRC and PKC, which have been shown to activate YAP1 directly through phosphorylation or indirectly through regulation of Rho GTPases and LATS activity (39–44). Of interest, it was recently reported that noncanonical WNT- and FZD8-mediated calcium signaling counteracts the tumorigenic activity of oncogenic KRAS (45). In contrast, FZD1 was shown to be important for WNT5A-mediated YAP1 activation (25). It is possible that the engagement of specific receptors by WNT5A determines its signaling and biological output in PDAC. Whether any one or all of these mechanisms are responsible for WNT5A-mediated YAP1 activation in PDAC remains to be further studied. Furthermore, additional noncanonical WNT ligands are also likely involved in YAP1 activation in PDAC. For instance, WNT7B expression is also elevated in the squamous subtype of PDAC and correlated with the YAP1 activation signature (data not shown). It is worth to determine whether the additional noncanonical WNT ligands also contribute to YAP1 activation in PDAC.

Although our data indicate that WNT5A overexpression in tumor cells functions in a cell-autonomous manner to activate YAP1 oncoprotein, tumor cells may also activate WNT5A signaling through paracrine mechanisms. Notably, WNT5A has been shown to be highly expressed in PDAC stroma fibroblast (46, 47), and our preliminary data suggest that the stromal WNT5A level is significantly correlated with tumor cell YAP1 level in human PDAC (data not shown). Therefore, stromal WNT5A could possibly contributes to YAP1 activation in tumor cells, given that the exuberant desmoplastic stroma is a defining characteristic of PDAC (2). In this scenario, the tumor-stroma interaction will thus play an instrumental role in orchestrating heterogeneous YAP1 activation in bulk tumor, which may in turn define the molecular heterogeneity and diverse biological phenotypes of PDAC.

Taken together, with agents targeting the Hippo-YAP pathway under development (30), our study showed that the critical role of WNT5A-mediated YAP1 activation in a subset of pancreatic tumors of the squamous subtype provides viable therapeutic targets for this most malignant form of human PDAC.

## Methods

### Transgenic mice

For the generation of a tamoxifen-inducible PDAC GEM model, *Mist1^Cre^*^ERT2/+^ (48) mice were used for conditional activation of mutant *Kras^G12D^* and mutant *Trp53^R172H^* in the mature pancreas. For *Yap1* deletion, these mice were further crossed with *Yap1^fl/fl^* mice (49). For the most efficient recombination, tamoxifen was administered intraperitoneally to 6-week-old mice in corn oil once daily for 5 days. The recombination efficiency was tested using PCR primers designed specifically to detect wild-type and recombinant alleles of *Kras*, *Trp53*, and *Yap1* in pancreatic tissues.

### Cell culture and establishment of primary PDAC lines

Human pancreatic cell lines SNU410, HPAC, HPAFII, PL45, PaTu8988S, and PaTu8988T were obtained from the American Type Culture Collection (ATCC). Pa04C was established from resected patient tumors, maintained as low passage (<10) (50), and cultured according to recommended protocols. Establishment and maintenance of primary mouse PDAC lines was performed as described previously (8, 14). Mouse PDAC cell line PD3077 was a gift from Dr. Ben Stanger, University of Pennsylvania Perelman School of Medicine. The human patient PDX cell lines were maintained in RPMI-1640 medium containing 10% FBS (Clontech). KRAS mutation status, molecular subtypes, and tumor grade information are listed in Supplementary Table 9.

### Reagents

Doxycycline (RPI), PE Annexin V Apoptosis Detection Kit I (BD Biosciences), BOX5 (EMD Millipore).

### Immunostaining and Western blot analysis

Immunohistochemical (IHC) analysis was performed as described previously (51). Details for Immunofluorescence staining, Western blot analysis, and primary antibody information are described in the Supplementary Methods.

### Lentivirus-mediated shRNA knockdown

All lentiviral shRNA clones targeting YAP1, WNT5A, and nontargeting shRNA control were obtained from Sigma Aldrich in the pLKO vector. The clone IDs for the shRNA are listed in the Supplementary Materials.

### CRISPR-Cas9–mediated gene knockout (KO)

SgRNAs targeting mouse *Wnt5a* or *Yap1* were cloned into pSpCas9(BB)-2A-Puro (Addgene, #62988) and transfected into target cells. After 2 µg/ml puromycin selection for 1 week, single cell clones were isolated and analyzed by the T7E1 assay and Western blot analysis. Sequences for *Wnt5a* and *Yap1* sgRNA are listed in Supplementary Materials.

### TMA staining and analysis

Immunohistochemical staining for YAP1 was performed on 5-µm unstained sections from the tissue microarray blocks, which included 92 (MD Anderson Cancer Center) or 83 (Johns Hopkins University School of Medicine) PDAC samples from patients who underwent upfront surgery. Immunohistochemical staining for YAP1 was reviewed by a pathologist (H.W.). The expression of YAP1 was classifies as YAP1-low and YAP1-high by using the median score for total YAP1 expression (nuclear plus cytoplasmic expression) as a cutoff.

### Statistical analysis

Tumor volume and tumor-free survival results were analyzed using GraphPad Prism. To assess distributional differences of variance across various test groups, the Mann-Whitney test was used. Other comparisons were performed by using the unpaired two-tailed Student *t* test. A *P* value less than 0.05 is considered significant. For all experiments with error bars, standard deviation (SD) was calculated to indicate the variation with each experiment and data, and values represented mean ± SD.

### Study approval

All animal studies were approved by MD Anderson Cancer Center Institutional Animal Care and Use Committee (IACUC) under protocol number 00001549.

## Author contributions

B.T., W.Y., S.F-B., S.C., Q.W., L.Y., X.Z., and S.G. performed data collection and interpretation. J.Y. provided statistical and bioinformatics analysis. C.Z. and S.B. contributed to animal breeding. Q.C., C.B., Y.K., H.Z., H.W., J.F., M.K. T.H. and A.M. provided clinical specimen and data analyses on PDX samples. J.Z., H.W., and A.M. performed pathology analyses. G.D., D.P., and A.M. provided crucial feedback on the manuscript. W.Y., S.G., and H.Y. drafted the manuscript. W.Y., S.G., and H.Y. conceived, designed, and supervised the study.

## Acknowledgments

We thank the laboratory of Dr. Ben Stanger for sharing mouse PDAC cell line PD3077. We would like to thank the Institute for Applied Cancer Science (IACS), the Flow Cytometry and Cellular Imaging Core at The University of Texas MD Anderson Cancer Center, and the Department of Veterinary Medicine at MD Anderson (Cancer Center Support Grant CA016672). We thank Dr. Ronald DePinho, Dr. Alan Wang, Dr. Mien-Chie Hung, Dr. Guocan Wang, Dr. Baoli Hu, Dr. Xin Zhou, and Dr. Jihye Paik for helpful discussions and critical reviews. The research was supported by the Pancreatic Cancer Action Network-AACR Career Development Award and NCI grant R01CA214793 to H.Y.; the Pancreatic Cancer Action Network-AACR Pathway to Leadership Award to W.Y.; Seed Grant from Hirshberg foundation for pancreatic cancer research to H.Y. and W.Y., and NCI P01 Grant P01CA117969 to H.W., J.B.F., M.K., G.F.D., A.M., and H.Y.

## Supplementary Materials and Method

### Immunostaining and Western Blot Analysis

For immunofluorescence staining, mouse and human cells were fixed with 4% paraformaldehyde-PBS for 15 min. Following Triton-X100 permeabilization and blocking, cells were incubated with primary antibodies overnight at 4°C following with Alexa 594-conjugated secondary antibodies at 4°C for 1 hour (Thermo Fisher Scientific, 1:1000). Samples were mounted using VECTASHIELD Antifade Mounting Medium with DAPI (Vector Laboratories) and immunofluorescence was detected using Olympus confocal microscopy. For western blot analysis, cells were lysed on ice using RIPA buffer supplemented with protease and phosphatase inhibitors (Sigma).

Primary Antibodies for Immunostaining and Western Blot Analysis: Yap (14074, Cell Signaling), pYAP (4911, Cell Signaling), Lats1 (3477, Cell Signaling), pLats1(8654, Cell Signaling), Wnt5a (2530, Cell Signaling), cleaved Caspase-3 (9661, Cell Signaling), Ki-67 (VP-K451, Vector Laboratories), Cyr61 (sc-13100, Santa Cruz Biotechnology), CTGF (sc-14939, Santa Cruz Biotechnology), AXL (8661, Cell Signaling), pErk (4376, Cell Signaling), pMEK (4376, Cell Signaling), Ck-19 (16858-1-AP, Proteintech), Actin (A2228, Sigma Aldrich), Vinculin (V4139, Sigma Aldrich), Kras (sc-30, Santa Cruz Biotechnology).

### Ectopic expression of YAP1 and WNT5A in mouse and human cells

To generate YAP1^S127A^-expressing stable Pa04C cells, Pa04C cells were transfected with a linearized pcDNA3.1 plasmid with or without YAP1 cDNA containing S127A substitution. Two days post-transfection using Lipofectamine1000, cultures were selected in G418 (Sigma) and single clones were picked and expanded for further analysis. Overexpression of YAPS127A or WNT5A in human or mouse cells other than Pa04C were acheieved with lentivral infection. Briefly, lentivirus infection was performed by transfecting 293T cells with either GFP control, YAP1^S127A^, or WNT5A cloned in pHAGE lentivirus vector {EF1α promoter-GW-IRES-eGFP (GW: Gateway modified)}. The virus was concentrated using ultracentrifuge and added to target cells in a 6-well plate containing 10ug/ml of polybrene (Millipore). 48 hours after infection GFP positive cells were selected by flow sorting.

### Lentivirus Mediated shRNA Knockdown

The clone IDs for shRNA are as follows: sh_mouse Yap1-1 (TRCN0000238436), sh mouse Yap1-2 (TRCN0000095864), sh_huYap1-1 (TRCN0000107265), sh_huYap1-2 (TRCN0000107266), sh_huWnt5a-1 (TRCN0000062717), sh_huWnt5a-2 (TRCN0000288987), sh_hu Kras-1 (TRCN0000033260), sh_hu Kras-2 (TRCN0000033262).

### Crispr-Cas9 Mediated Gene Knockout

Sequences for *Wnt5a* sgRNA are as follows: sgRNA-1 F: CTTGAGAAAGTCCTGCCAGT; R: ACTGGCAGGACTTTCTCAAG. sgRNA-2 F: GAAACTCTGCCACTTGTATC; R: GATACAAGTGGCAGAGTTTC. sgRNA-3 F: TATACTTCTGACATCTGAAC; R: GTTCAGATGTCAGAAGTATA. sgRNA-4 F: ACAGCCTCTCTGCAGCCAAC, R: GTTGGCTGCAGAGAGGCTGT.

Sequences for *Yap1* sgRNA are as follows: sgRNA-1 F: ACCAGGTCGTGCACGTCCGC; R: GCGGACGTGCACGACCTGGT. sgRNA-2 F: CCCCGCGGACGTGCACGACC; R: GGTCGTGCACGTCCGCGGGG.

### Xenograft Studies

For orthotopic xenografts, 5 × 10^5^ cells were injected pancreatically into NCr nude mice (Taconic) and tumor growth was monitored with bioluminescent imaging as described^1^. For Sub-Q xenografts, 1 × 10^6^ cells (mouse tumor cells) or 3 × 10^6^ cells (human PDAC or PDX cells) were injected subcutaneously into the lower flank of NCr nude mice. Tumor volumes were measured every 7 days starting from Day 7 post injection and calculated using the formula, volume = length × width^2^/2.

### Immunoprecipitation Assay

Immunoprecipitation of YAP1 complexes was performed by lysing cells with 1% NP40 lysis buffer containing phosphatase and protease inhibitor cocktails on ice for 45 minutes. 1 mg of lysate was incubated overnight at 4°C with primary antibodies followed with protein A/G Plus-agarose (sc-2003, Santa Cruz) for 3 hours at 4^ο^C. Immunoprecipitates were washed three times with lysis buffer, then re-suspended with 2X sample buffer boiled for 5 min and detected by western blot analyses.

### Quantitative Real-time Polymerase Chain Reaction Analysis

RNA from cell lines and pancreas tissues was isolated using RNeasy Mini Kit (Qiagen) and first-strand cDNA was synthesized from 2 µg total RNA using random primers and Omniscript® Reverse Transkriptase Kit (Qiagen). Actin was used as housekeeping gene. Real-time polymerase chain reaction experiments were performed in triplicates and are displayed ± SD.

### Clonogenic assay

500-2000 cells were seeded into each well of 6-well plate in triplicates and incubated to allow colony formation for 10-20 days. The colonies were stained with 0.2% crystal violet in 80% methanol for 30 minutes at room temperature and de-stained upon which they were scanned and colonies counted using Image J (http://rsb.info.nih.gov/ij/).

### Gene Expression and PDAC-Subtype Analysis

mRNA expression profiling on Illumina microarrays were performed according to the manufacture’s protocol. Raw data was processed using Genome Studio (GSGX Version 1.6.0) and analysis was done using group quantile normalization with background subtraction. Complete profiles are available at GEO at GSE135754. The software package LIMMA (Linear Models for Microarray Data) was applied to detect significantly differentially expressed probes using Benjamini-Hochberg adjusted p-values. For GSEA analysis, gene sets collection from MSigDB 3.0 and Kyoto Encyclopedia of Genes and Genomes (KEGG) were included in the analysis.

For molecular subtype analysis, we combined subtype specific genes from Collisson et al and Bailey et al studies to construct a “NanoString signature” comprised of 32, 23, and 17 subtype specific genes for squamous, progenitor, and ADEX subtypes, respectively (see supplementary table 2). To call PDAC subtypes in human PDAC cell lines or PDXs, we used the following algorithm. First, subtype signature scores were calculated by summing up Z scores from subtype specific genes which were ceiled at 2.5 and bottomed at −0.5. Kmeans two separation were then done to call high and low groups within each subtype. SQ/Pro subtypes are then called as squamous gene high/progenitor gene low and vice versa. ADEX subtype is called as ADEX gene high and SQ/Pro gene low. The remaining tumors are undefined. The nanoString call results are further confirmed by heatmap visualiation using Collission genes (data not shown)^2^. The clusters were viewed using Java TreeView (version 1.1.6r4) ^3^.

---

**Supplementary Figure 1.**
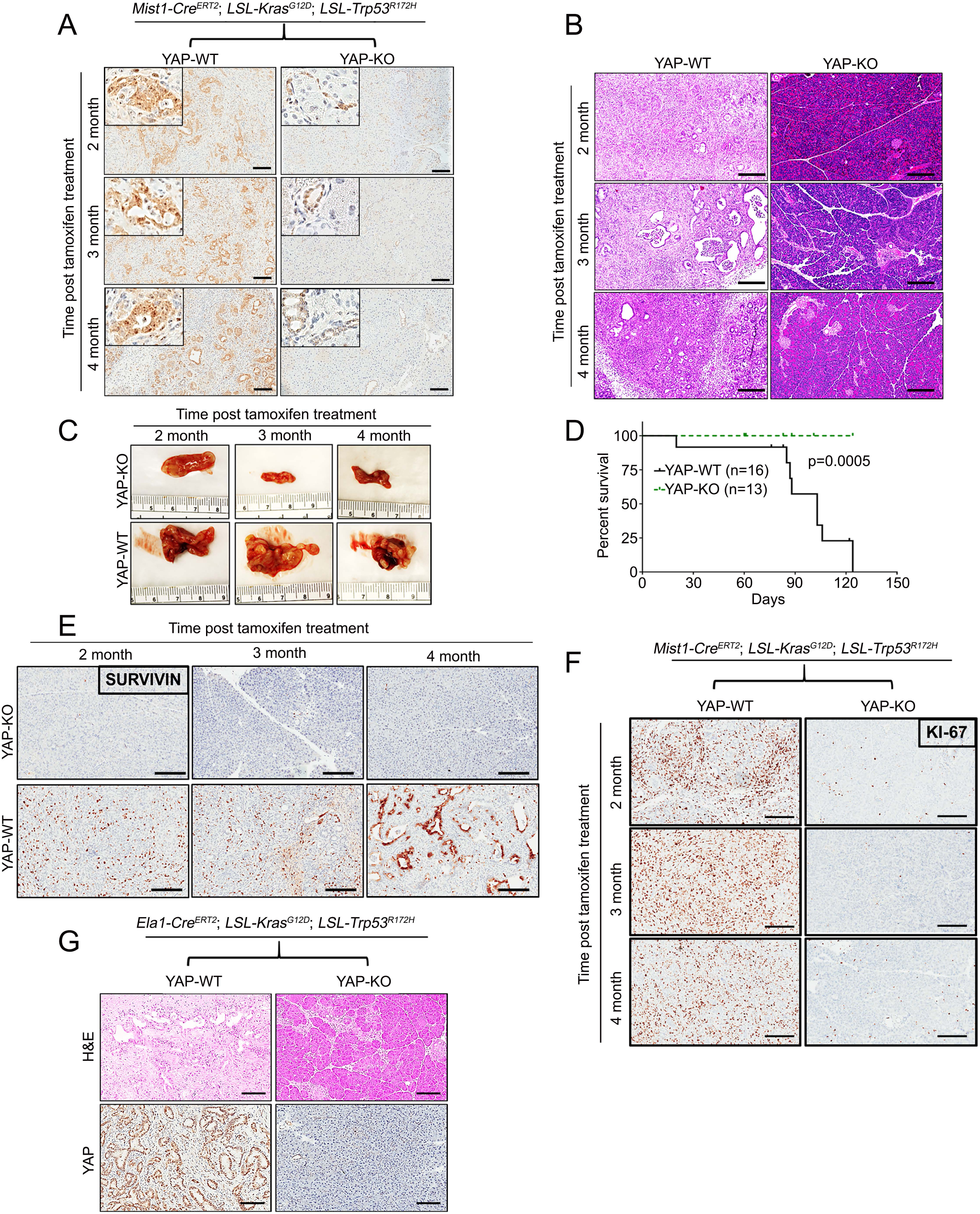
YAP1 is essential for PDAC development. (A) *Yap1* wild-type (WT) or knockout (KO) MKP mice driven by tamoxifen-in­ ducible Mist1-CreERT2. Pancreata were collected at indicated time points, and YAP1 staining shows YAP1 activation in PDAC tumors of MKP mice. Scale bar: 50 µm. (B-C) Gross images of pancreata (B) and H&E staining (C) show block of tumor development in *Yap1-KO* mice. Scale bar: 100 µm. **(D)** Kaplan-Meier overall survival analysis for *Yap1-WT* and *Yap1-KO* MKP mice. The *P* value for survival analysis was calculated with the log rank test. **(E-F)** Pancreata from *Yap1* WT or KO MKP mice were collected at indicated time points and stained for Survivin **(E)** or Ki67 **(F).** For each time point (age group), at least 4 mice per genotype were analyzed. Scale bar: 100 µm. **(G)** *Yap1* wild-type (YAP-WT) or knockout (YAP-KO) *Kras^G12D^;Trp53^R172H^* mice driven by *Elastase-Cre^ERT2^*upon tamoxifen injection. Pacreata collected at 4 months post-injection and stained for H&E (Top) and YAP1 (Bottom) Scale bar: 100µm.

**Supplementary Figure 2.**
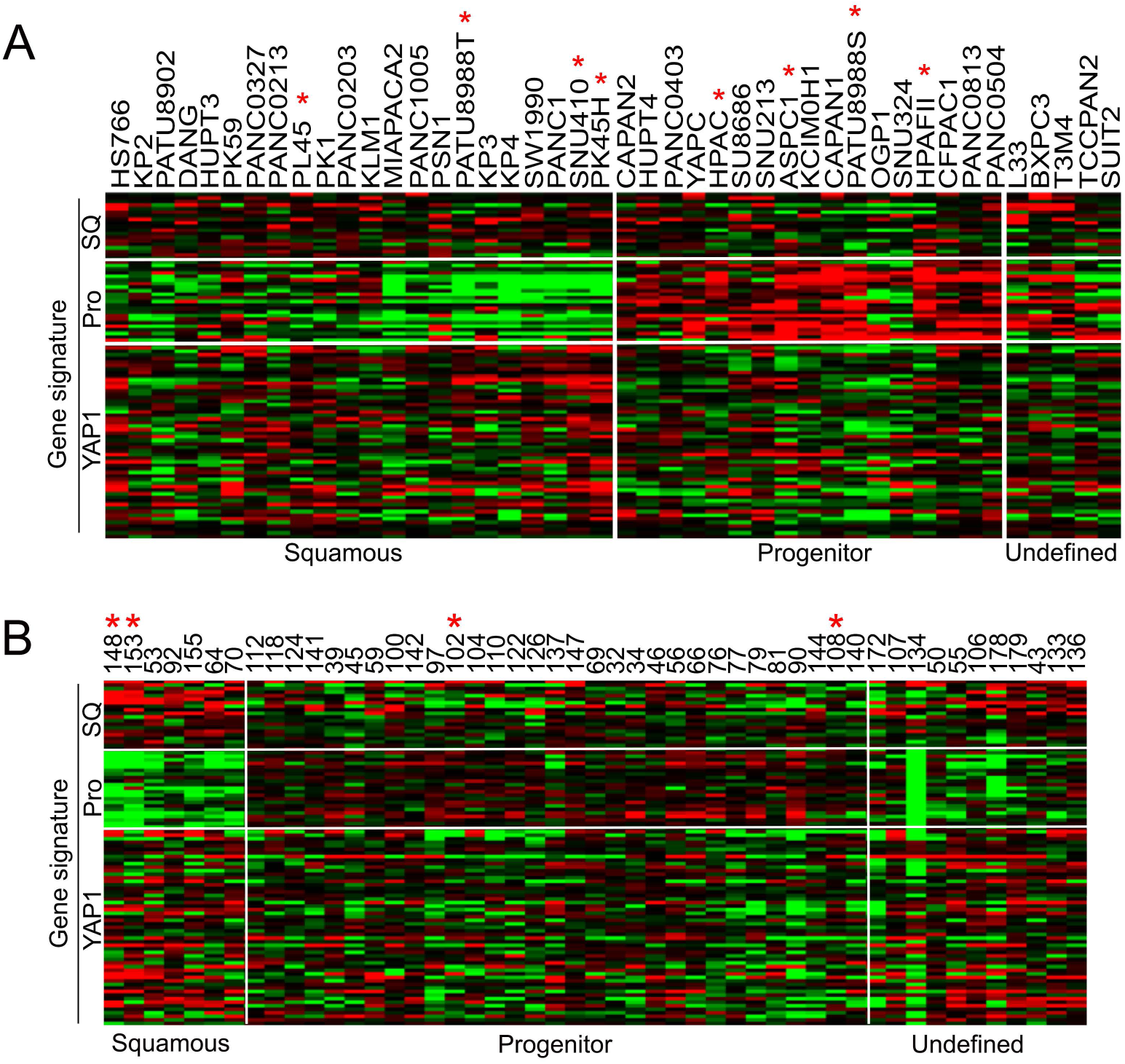
YAP1 is activated in the squamous subtype of human PDAC. **(A-B)** Clustering of human PDAC cell lines **(A)** and human PDXs **(B)** based on subtype-specific gene signatures. Heatmap of squamous (SQ), progenitor (PRO), and YAP1 activation signature is shown. Asterisk: cell lines used in the stydy.

**Supplementary Figure 3.**
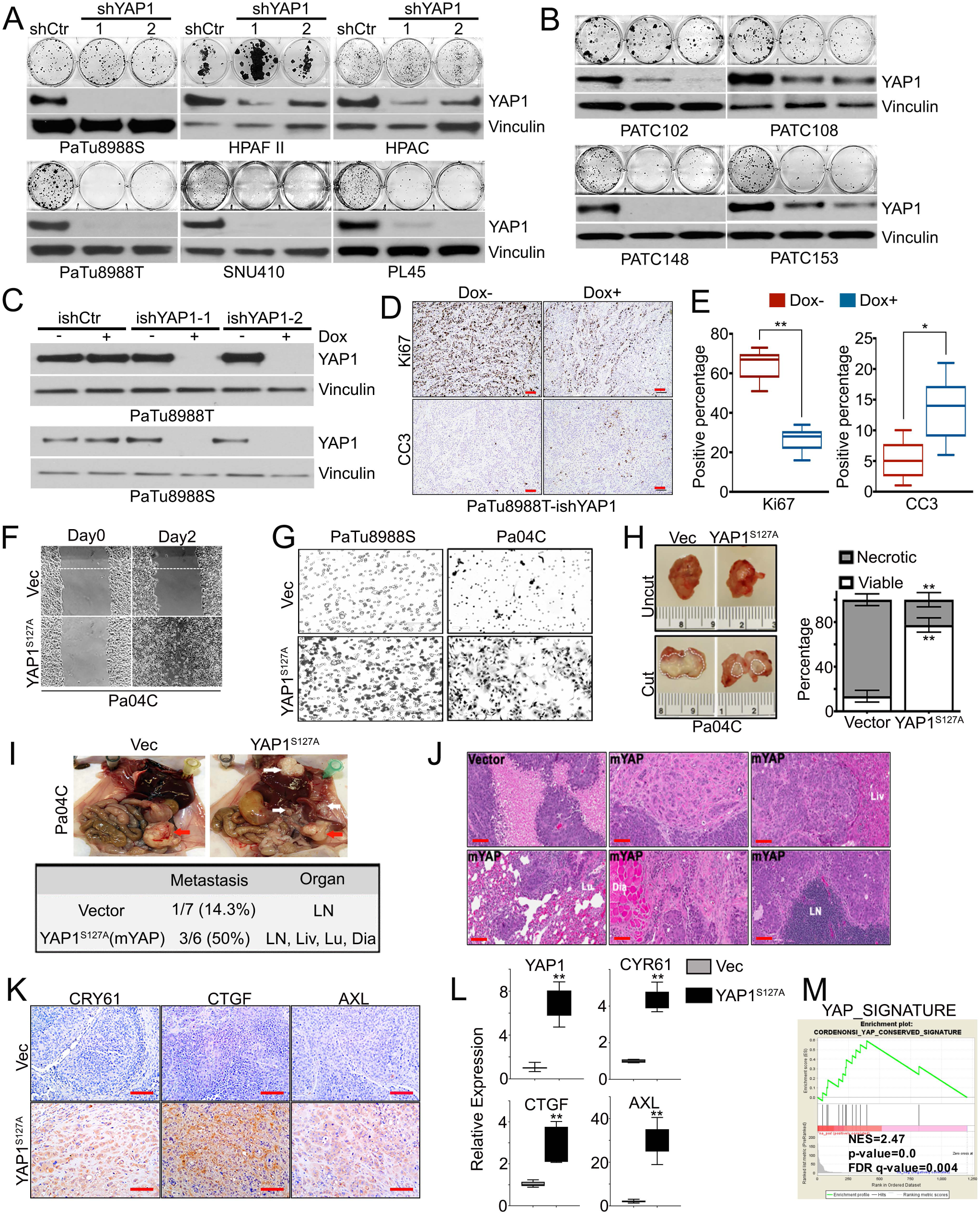
YAP1 is essential for the maintenance of squamous subtype of PDACs. **(A-B)** Representative images of the colony formation assay in human PDAC cell lines **(A)** or POX cells **(B)** infected with YAP1 shRNAs or non-targeting shRNA (shCtr). The YAP1 knockdown efficiency was detected by Western blot. **(C)** Western blot analysis for YAP1 in PaTu8988S and PaTu8988T cells engineered with inducible shRNA targeting YAP1 in the presence or absence of doxycycline for 72 hours. Inducible non-targeting shRNA (ishCtr) was used as control. **(D)** Subcutaneous xenograft tumors from inducible shYAP1-infected PaTu8988T cells were treated with vehicle or doxycycline for 1 week. Tumors were collected and stained for Ki67 or cleaved Caspase3 (CC3). Scale bar: 100 µm. Percentage of positive cells is shown in **(E).** Error bars represent ±SD (n=6 fields, 250 cells/field). *: p<0.05; **: p<0.01. **(F)** In vitro wound healing assay for Pa04C cells expressing YAP1^S127A^ or control vector (Vee). **(G)** Ectopic expression of YAP1^S127A^ in PaTu8988S and Pa04C cells promotes cell invasion in a Boyden chamber assay. **(H)** Representative gross images of orthotopic tumors from Pa04C-Vector (Vee) or Pa04C-YAP^S127A^ cells. Left panel shows uncut tumors (top) and tumors cut in half to reveal necrotic area (bottom). Right panel shows quantification of necrotic area. **: *P* < 0.01. **(I)** Orthotopic xenograft tumors from Pa04C-Vector or -YAP1^S127A^ cells (top). Red arrows indicate primary tumors and white arrows indicate metastatic tumors grown on peritoneum, lymph node and liver. Metastasis rate and involved organs are summarized in bottom panel. **(J) H&E** staining of orthotopic primary and metastatic tumors show metastases of Pa04C-YAP^S127A^ (mYAP) cells in liver (Liv), lung (Lu), diaphragm (Dia) and lymph node **(LN).** Scale bar: 1OOµ m. **(K)** Orthotopic xenograft tumors from Pa04C cells expressing vector (Vee) or YAP1^S127A^ were stained for CYR61, CTGF and AXL. Scale bar: 1OOµm. **(L)** Relative mRNA levels of YAP1 and indicated YAP1-target genes in Pa04C cells expressing vector (Vee) or YAP1^S127A^. Error bars indicate ±SD of triplicates. **: *P* < 0.01. **(M)** GSEA plot of YAP1 activation signature based on the gene expression profiles of vector-vs YAP1s 1_27A e xpressing Pa04C cells. NES denotes normalized enrichment score.

**Supplementary Figure 4.**
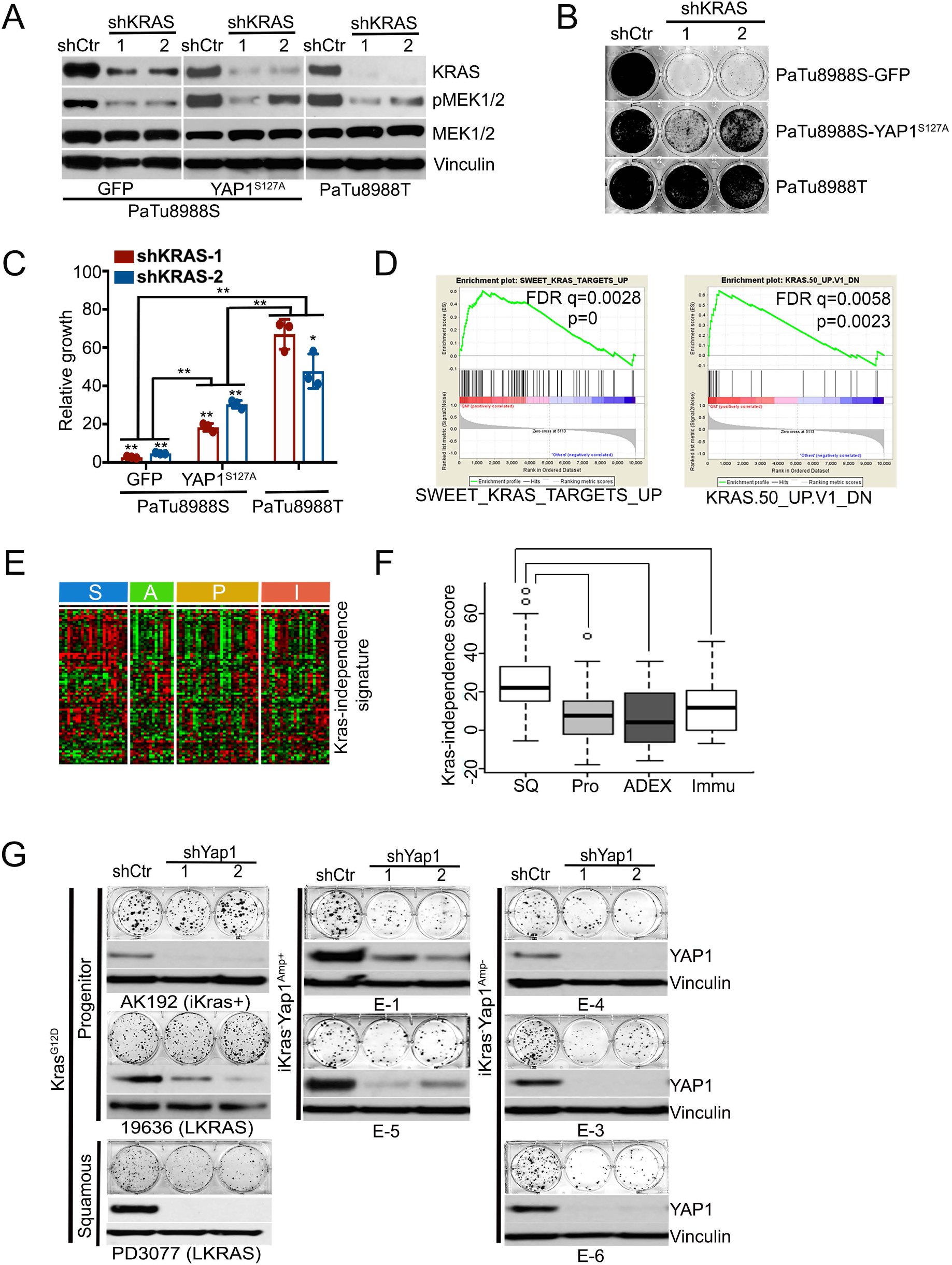
YAP1 bypasses KRAS-dependency in PDAC. **(A)** Western blot analysis for KRAS, phosphor-MEK1/2 and MEK1/2 in PaTu8988S cells expressing GFP or YAP15127A and PaTu8988T cells upon knockdown of KRAS with two independent shRNAs. Non-targeting shRNA (shCtr) was used as control. **(B)** Representative images of the cell growth assay in human PDAC cell lines infected with KRAS shRNAs or non-targeting shRNA (shCtr). Quantification from triplicates is shown in **(C)** and is presented as relative growth upon normalization to shCtr group. Error bars indicate ±SD. *: P< 0.05; **: *P* < 0.01. **(D)** GSEA plots show genes induced upon KRAS knockdown or suppressed upon KRAS expression are enriched in squamous subtype human PDAC. **(E)** Heatmap shows the expression of KRAS-independence gene signature among human PDAC subtypes in PDAC TCGA dataset with quantification shown in **(F).** S: squamous subtype; A: ADEX subtype; P: progenitor subtype; I: immunogenic subtype. **(G)** Representative images of the colony formation assay in mouse PDAC cells infected with Yap1 shRNAs or non-targeting shRNA (shCtr). The YAP1 knockdown efficiency was detected by Western blot.

**Supplementary Figure 5.**
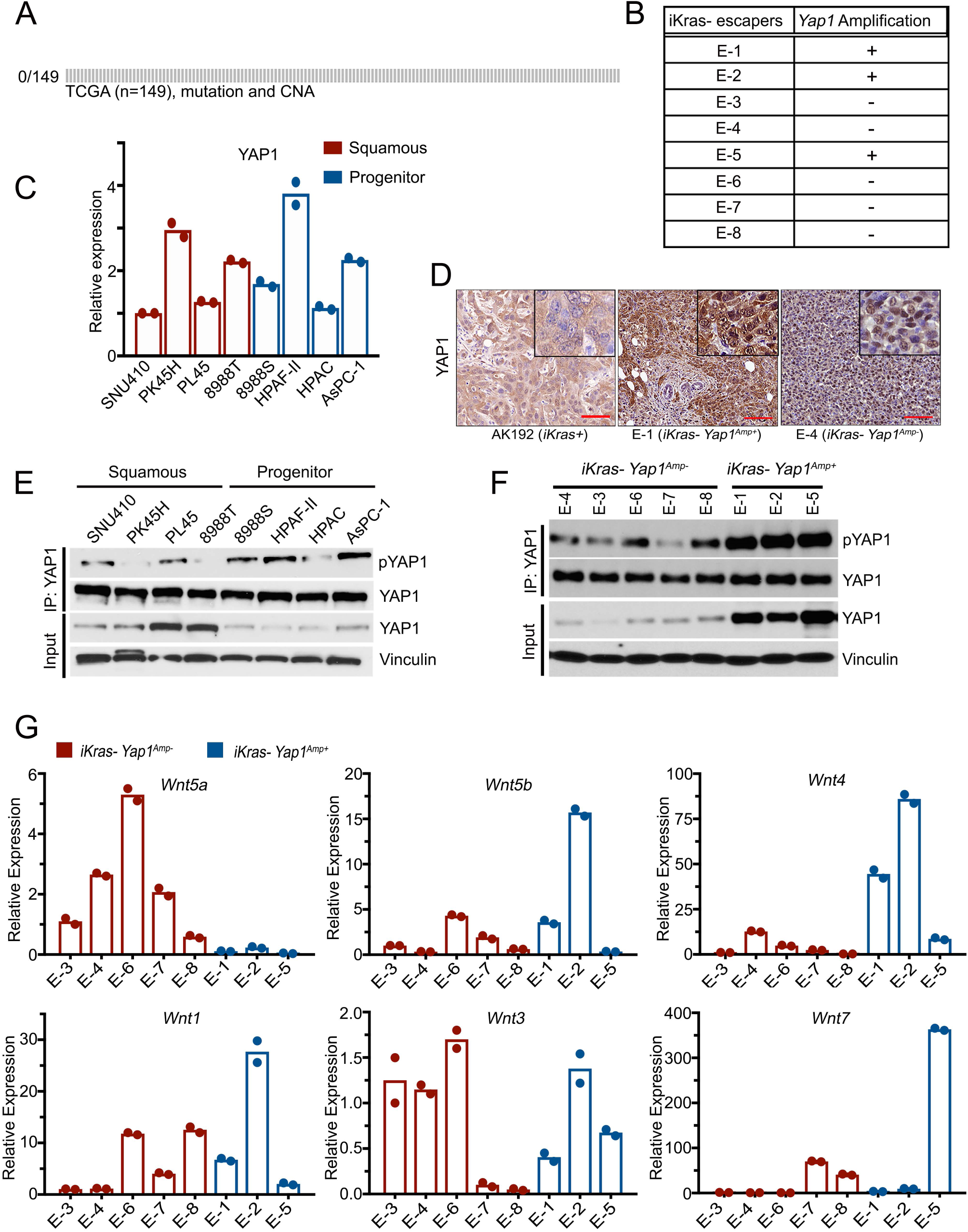
YAP1 activation in PDAC is mediated by WNT5A overexpression. **(A)** Mutation and copy number alteration (CAN) re­ quency of *YAP1* in human PDAC from TCGA datasets. **(B)** *Yap1* gene amplification status in *iKras-* tumors. **(C)** mRNA expression level of YAP1 in human PDAC cell lines (n=2). **(D)** *iKras+, iKras-Yap1^Amp-^* and *iKras-Yap1^Amp+^* tumors were stained for YAP1. **(E-F)** Equal amount of YAP1 protein was immunoprecipitated from human PDAC cell lines **(E)** or mouse iKras-PDAC cells **(F)** and subjected to western blot analysis for phosphor-YAP1 (S127) and YAP1 (top). Input shows the western blot analysis for YAP1 in whole cell lysates (bottom). **(G)** Relative mRNA levels of indicated WNT ligands in mouse iKras-PDAC cells (n=2).

**Supplementary Figure 6.**
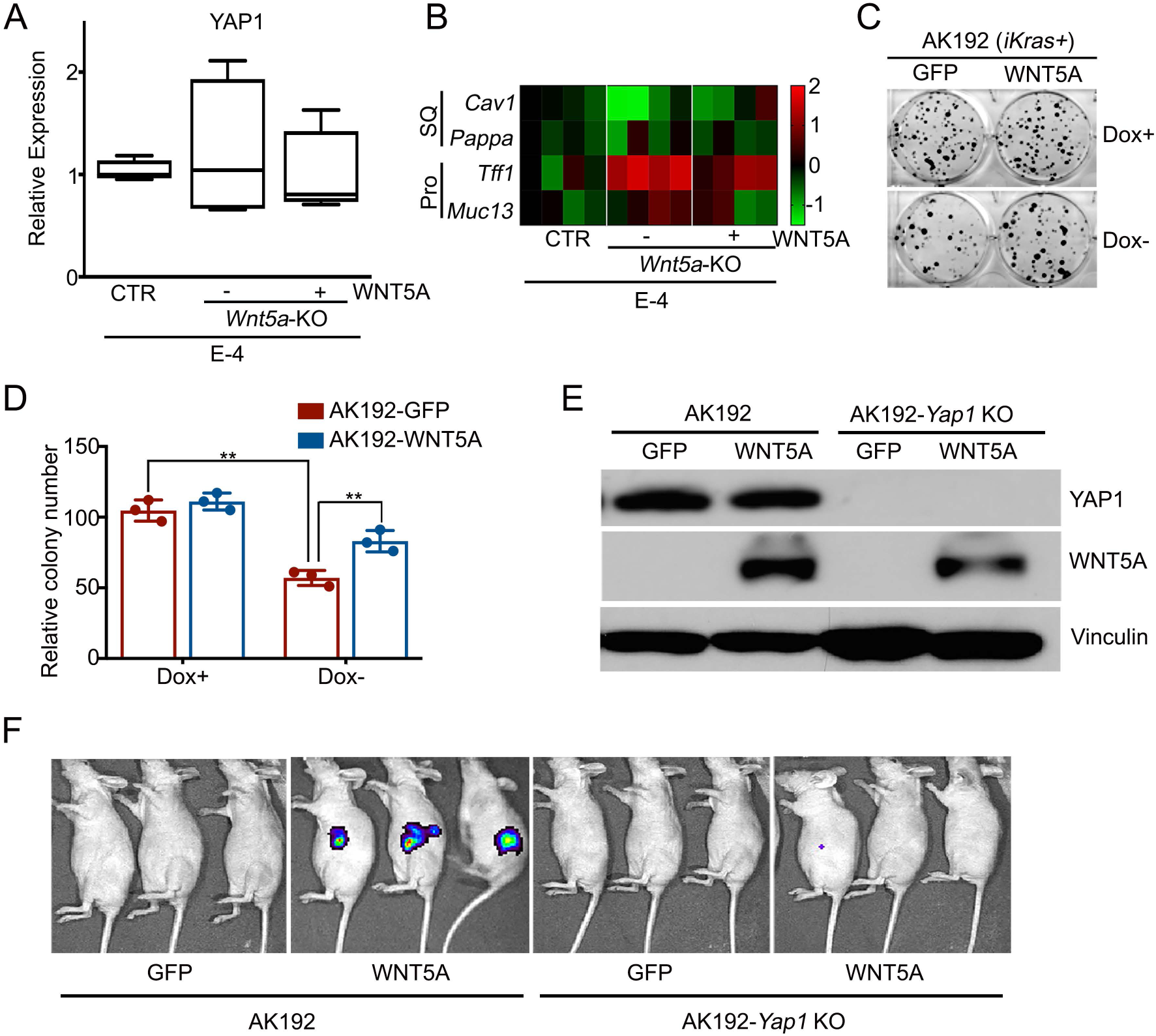
WNT5A overexpression enables tumor maintenance and bypass of KRAS dependence. **(A)** Expression of Yap1 or subtype-specific genes **(B)** in subcutaneous xenograft tumors generated with E-4 *(iKras-Yap1^Amp-^)* and *E4-Wnt5a* KO cells infected with GFP or WNT5A. **(C)** Representative images of the colony formation assay in mouse AK192 *(iKras+)* cells infected with GFP or WNT5A grown in the presence (Dox+) or absence (Dox-) of doxycycline. Quantification of colony formation from triplicates is shown in **(D).** Error bars from all panels indicate ±SD. *: *P* < 0.05. **(E)** Western blot analysis for YAP1 and WNT5A in AK192 or AK192-Yap1 KO cells infected with GFP or WNT5A. **(F)** AK192 or AK192-Yap1 KO cells expressing luciferase were infected with lentivirus expressing GFP or WNT5A and orthotopically injected into nude mice pancreas in the presence of doxycycline. Animals were withdrawn from doxycycline four days later and tumor growth was visualized by bioluminescent imaging at 6 weeks.

**Supplementary Table 1.**
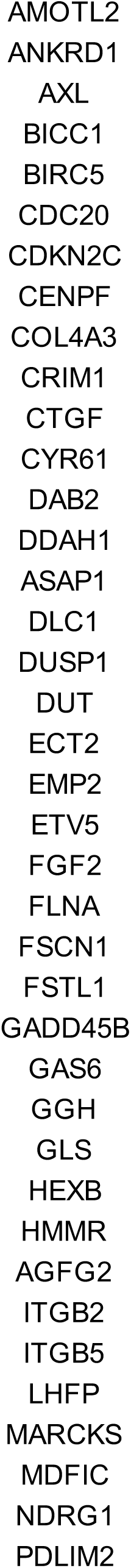
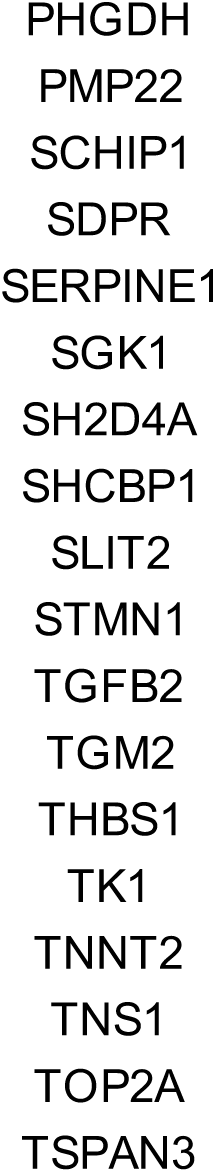
YAP1 signature: CORDENONSI_YAP_CONSERVED_SIGNATURE (GSEA MSigDB database)

**Supplementary Table 2.** Subtype signature genes used for NanoString analysis based on Collisson et al and Bailey et al studies (Nat Med. 2011, 17:500-3, Nature. 2016, 531:47-52)

| **Gene** | **Subtype** | **<u>Source:</u>**<br>**Collisson** | **<u>Source:</u>**<br>**Bailey** |
| --- | --- | --- | --- |
| TNC | Squamous | 0 | 1 |
| PTHLH | Squamous | 0 | 1 |
| TMEM40 | Squamous | 0 | 1 |
| KLC3 | Squamous | 0 | 1 |
| CAV1 | Squamous | 1 | 1 |
| TGFBI | Squamous | 0 | 1 |
| GJB6 | Squamous | 0 | 1 |
| CDH26 | Squamous | 0 | 1 |
| EREG | Squamous | 0 | 1 |
| RHCG | Squamous | 0 | 1 |
| RGS20 | Squamous | 0 | 1 |
| GSDMC | Squamous | 0 | 1 |
| HMGA2 | Squamous | 0 | 1 |
| C16orf74 | Squamous | 0 | 1 |
| IVL | Squamous | 0 | 1 |
| A2ML1 | Squamous | 0 | 1 |
| NIPAL4 | Squamous | 0 | 1 |
| KRT5 | Squamous | 0 | 1 |
| KRT14 | Squamous | 1 | 1 |
| RFX8 | Squamous | 0 | 1 |
| S100A2 | Squamous | 1 | 1 |
| L1CAM | Squamous | 0 | 1 |
| KRT6A | Squamous | 0 | 1 |
| MIR205HG | Squamous | 0 | 1 |
| TWIST1 | Squamous | 1 | 0 |
| NT5E | Squamous | 1 | 0 |
| PMAIP1 | Squamous | 1 | 0 |
| LOX | Squamous | 1 | 0 |
| SLC2A3 | Squamous | 1 | 0 |
| GPM6B | Squamous | 1 | 0 |
| PAPPA | Squamous | 1 | 0 |
| AIM2 | Squamous | 1 | 0 |
| TFF1 | Progenitor | 1 | 1 |
| CEACAM5 | Progenitor | 1 | 0 |
| TFF3 | Progenitor | 1 | 0 |
| ATP10B | Progenitor | 1 | 1 |
| CAPN8 | Progenitor | 1 | 1 |
| CEACAM6 | Progenitor | 1 | 0 |
| TSPAN8 | Progenitor | 1 | 1 |
| TOX3 | Progenitor | 1 | 1 |
| GPX2 | Progenitor | 1 | 1 |
| SDR16C5 | Progenitor | 1 | 0 |
| PHGR1 | Progenitor | 0 | 1 |
| DPCR1 | Progenitor | 0 | 1 |
| SEMG1 | Progenitor | 0 | 1 |
| SLC9A3 | Progenitor | 0 | 1 |
| SULT1B1 | Progenitor | 0 | 1 |
| FAM177B | Progenitor | 0 | 1 |
| MUC17 | Progenitor | 0 | 1 |
| CAPN9 | Progenitor | 0 | 1 |
| EDN3 | Progenitor | 0 | 1 |
| CLDN18 | Progenitor | 0 | 1 |
| MYO7B | Progenitor | 0 | 1 |
| BTNL8 | Progenitor | 0 | 1 |
| IHH | Progenitor | 0 | 1 |
| CELA3A | ADEX | 1 | 1 |
| CTRB2 | ADEX | 1 | 1 |
| PLA2G1B | ADEX | 1 | 1 |
| REG1B | ADEX | 1 | 1 |
| CELA3B | ADEX | 1 | 1 |
| CLPS | ADEX | 1 | 1 |
| CEL | ADEX | 1 | 1 |
| PRSS1 | ADEX | 1 | 1 |
| GP2 | ADEX | 1 | 1 |
| CTRB1 | ADEX | 0 | 1 |
| SYCN | ADEX | 0 | 1 |
| CPA2 | ADEX | 0 | 1 |
| CPA1 | ADEX | 0 | 1 |
| REG3G | ADEX | 0 | 1 |
| CTRC | ADEX | 0 | 1 |
| AQP8 | ADEX | 0 | 1 |
| CUZD1 | ADEX | 0 | 1 |

**Supplementary Table 3.**
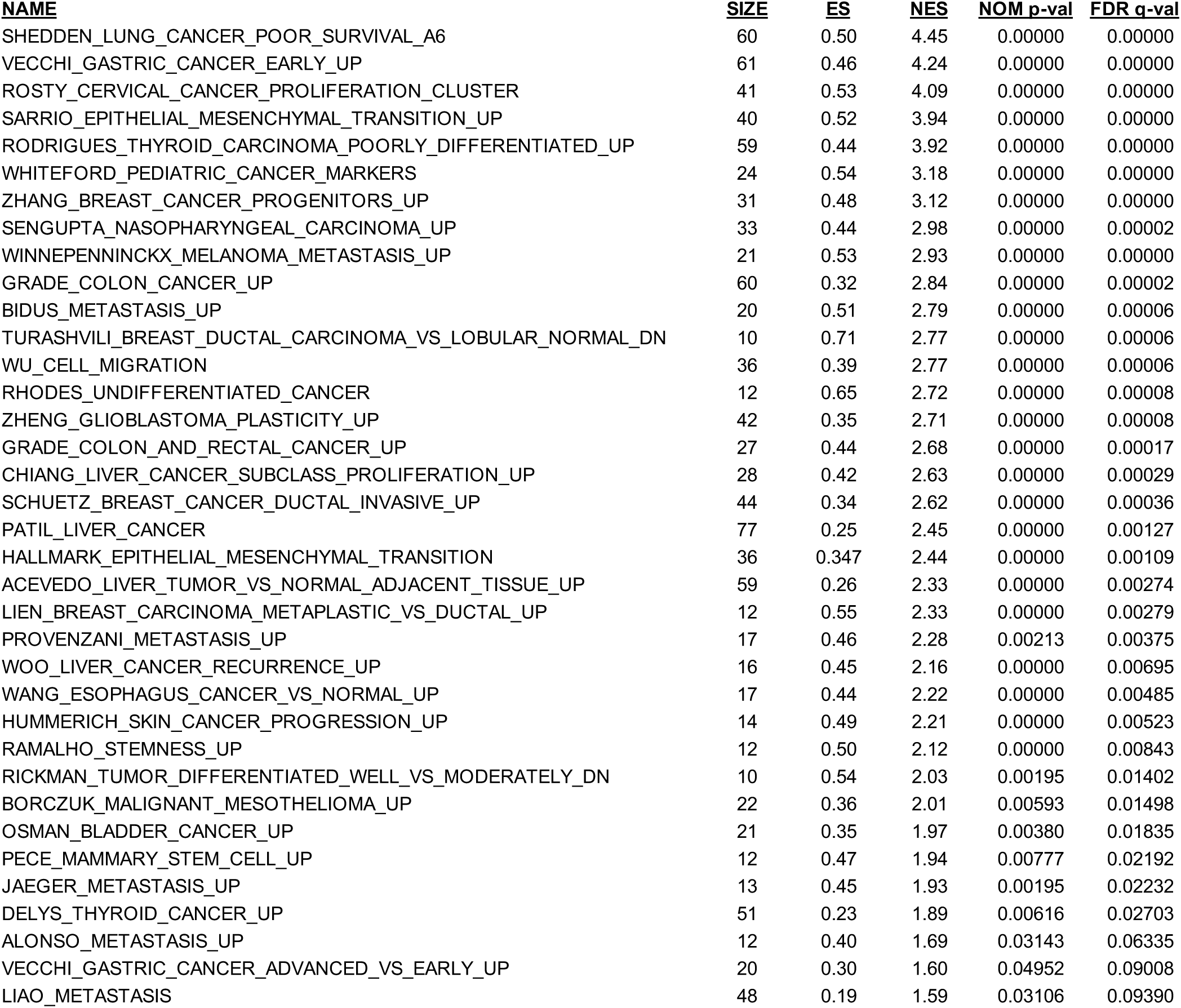
YAP1-regulated genes are enirched in signatures for tumor development and metastasis.

**Supplementary Table 4.**
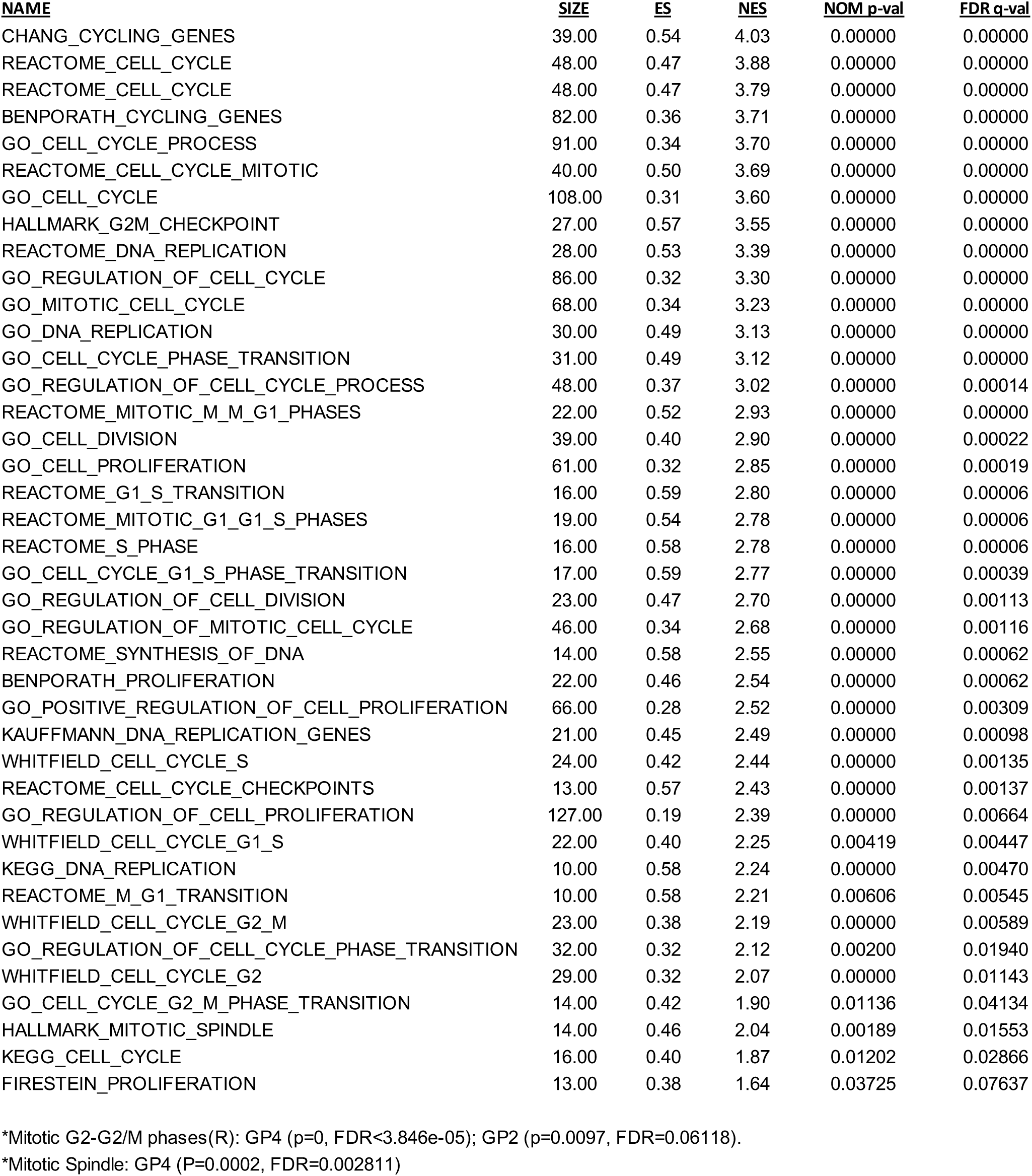
YAP1-regulated genes are enriched in signatures related to cell proliferation and cell cycle progression, part of GP2 and GP4 gene programs associated with squamous subtype of PDAC* (Bailey et al).

**Supplementary Table 5.** YAP1 targets are enriched in gene signatures associated with well known oncogenic signaling pathways and gene programs associated with squamous subtype of PDAC* (Bailey et al).

**Validated targets of C-MYC transcriptional activation(N)\***
| <u>NAME</u> | <u>SIZE</u> | <u>ES</u> | <u>NES</u> | <u>NOM p-val</u> | <u>FDR q-val</u> |
| --- | --- | --- | --- | --- | --- |
| HALLMARK_TNFA_SIGNALING_VIA_NFKB | 42 | 0.42 | 3.16 | 0.00000 | 0.00000 |
| HALLMARK_MYC_TARGETS_V2 | 10 | 0.67 | 2.57 | 0.00000 | 0.00000 |
| HALLMARK_MYC_TARGETS_V1 | 12 | 0.57 | 2.41 | 0.00000 | 0.00091 |
| MYC_UP.V1_UP | 14 | 0.45 | 2.06 | 0.00000 | 0.01826 |
| DANG_BOUND_BY_MYC | 71 | 0.28 | 2.69 | 0.00000 | 0.00017 |
| DANG_MYC_TARGETS_UP | 10 | 0.61 | 2.37 | 0.00000 | 0.00220 |
| BILD_MYC_ONCOGENIC_SIGNATURE | 23 | 0.39 | 2.30 | 0.00411 | 0.00350 |
| ALFANO_MYC_TARGETS | 24 | 0.32 | 1.82 | 0.01613 | 0.03544 |

\*GP5 (P=0, FDR<1.000e-04), GP4 (P=0.0021, FDR=0.01989), GP2 (P=0.016, FDR=0.08453)

**IL6-mediated signaling events(N)\***
| <u>NAME</u> | <u>SIZE</u> | <u>ES</u> | <u>NES</u> | <u>NOM p-val</u> | <u>FDR q-val</u> |
| --- | --- | --- | --- | --- | --- |
| CROONQUIST_IL6_DEPRIVATION_DN | 23 | 0.64 | 3.60 | 0.00000 | 0.00000 |
| DAUER_STAT3_TARGETS_UP | 11 | 0.50 | 2.07 | 0.00382 | 0.01121 |
| HALLMARK_IL6_JAK_STAT3_SIGNALING | 11 | 0.39 | 1.62 | 0.04211 | 0.07865 |

\*GP2 (P=0.0157, FDR=0.0835)

**E2F transcription factor network(N)\***
| <u>NAME</u> | <u>SIZE</u> | <u>ES</u> | <u>NES</u> | <u>NOM p-val</u> | <u>FDR q-val</u> |
| --- | --- | --- | --- | --- | --- |
| HALLMARK_E2F_TARGETS | 39 | 0.54 | 3.93 | 0.00000 | 0.00000 |
| E2F1_UP.V1_UP | 21 | 0.54 | 2.94 | 0.00000 | 0.00075 |
| ISHIDA_E2F_TARGETS | 13 | 0.58 | 2.56 | 0.00000 | 0.00058 |
| PID_E2F_PATHWAY | 14 | 0.51 | 2.33 | 0.00199 | 0.00275 |
| BILD_E2F3_ONCOGENIC_SIGNATURE | 32 | 0.31 | 2.08 | 0.00000 | 0.01104 |

\*GP4 (P=0; FDR<4.762e-05)

**RhoA signaling pathway(N)\***
| <u>NAME</u> | <u>SIZE</u> | <u>ES</u> | <u>NES</u> | <u>NOM p-val</u> | <u>FDR q-val</u> |
| --- | --- | --- | --- | --- | --- |
| BERENJENO_TRANSFORMED_BY_RHOA_UP | 82 | 0.47 | 4.81 | 0.00000 | 0.00000 |

\*GP3 (P=0.0029, FDR=0.02324)

**TGFB/BMP Receptor Signaling\***
| <u>NAME</u> | <u>SIZE</u> | <u>ES</u> | <u>NES</u> | <u>NOM p-val</u> | <u>FDR q-val</u> |
| --- | --- | --- | --- | --- | --- |
| LEE_BMP2_TARGETS_DN | 90 | 0.49 | 5.27 | 0.00000 | 0.00000 |
| TGFB_UP.V1_UP | 23 | 0.42 | 2.39 | 0.00000 | 0.00526 |
| COULOUARN_TEMPORAL_TGFB1_SIGNATURE_UP | 13 | 0.53 | 2.29 | 0.00000 | 0.00361 |
| KARLSSON_TGFB1_TARGETS_UP | 10 | 0.58 | 2.19 | 0.00199 | 0.00587 |
| PLASARI_TGFB1_TARGETS_10HR_UP | 36 | 0.29 | 1.93 | 0.00853 | 0.02192 |
| MCBRYAN_PUBERTAL_TGFB1_TARGETS_UP | 19 | 0.33 | 1.69 | 0.02863 | 0.06292 |

\*BMP receptor signaling(N):GP3 (P=0.0001, FDR=0.001106)

| <u>NAME</u> | <u>SIZE</u> | <u>ES</u> | <u>NES</u> | <u>NOM p-val</u> | <u>FDR q-val</u> |
| --- | --- | --- | --- | --- | --- |
| HALLMARK_TNFA_SIGNALING_VIA_NFKB | 42 | 0.42 | 3.16 | 0.00000 | 0.00000 |
| PHONG_TNF_TARGETS_UP | 15 | 0.65 | 2.99 | 0.00000 | 0.00000 |

\*GP2 (P= 0.0004; FDR=0.008161)

**Supplementary Table 6.** Yap1 targets in this study are enriched in Hippo signaling pathway (GP3) of squamous subtype (Bailey et al)

| <u>Bailey et al.</u> | <u>This study</u> |
| --- | --- |
| Hippo signaling pathway(K) GP3_Reactome (p=0, FDR<7.692e-05) | Pa04C-YAP<sup>S127A</sup> (Fold Change) |
| FZD10 | 128.00 |
| WTIP | 20.22 |
| FGF10 | 12.90 |
| TCF7L1 | 10.00 |
| TGFB1 | 9.70 |
| BMP8A | 7.06 |
| TGFBR1 | 6.80 |
| BMP6 | 5.81 |
| SMAD7 | 5.57 |
| CTGF | 3.28 |
| TGFB3 | 2.80 |
| LATS2 | 2.17 |
| SNAI2 | 2.05 |
| TEAD1 | 1.71 |
| GLI2 | 0.49 |
| FZD2 | 0.43 |
| LEF1 | 0.10 |
| DLG4 | 0.08 |

**Supplementary Table 7.**
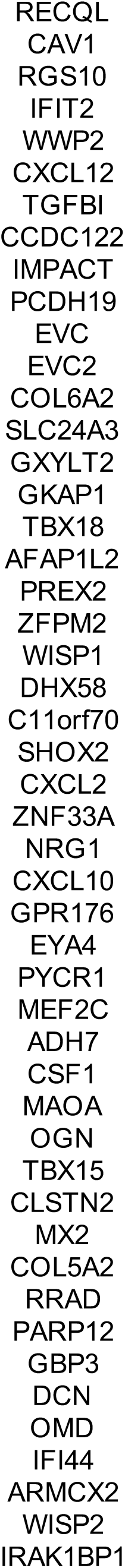
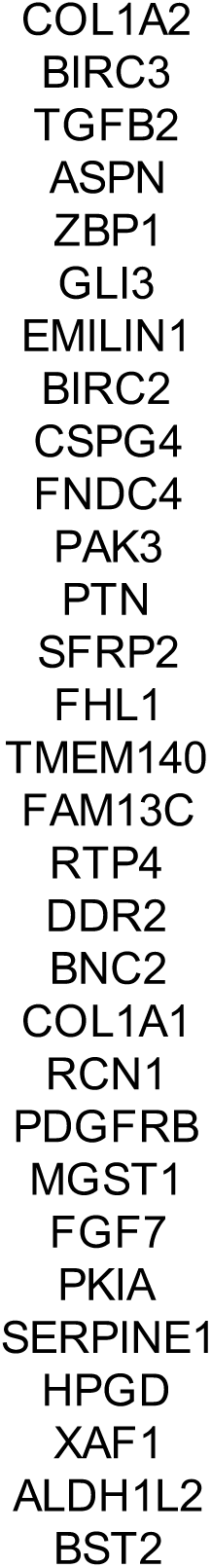
KRAS-independent signature based on Kapoor et al, Cell. 2014. 158 (1):185-197

**Supplementary Table 8.**
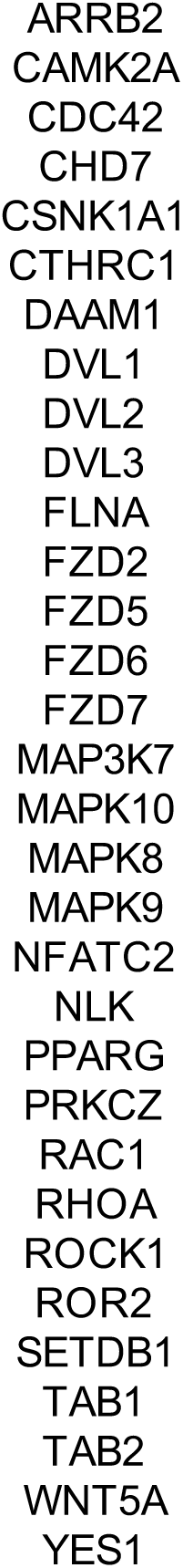
Non-canonical WNT signature: PID_WNT_NONCANONICAL_PATHWAY (GSEA MSigDB database)

**Supplementary Table 9.** Sample information for human PDAC cell lines and PDX lines

| Name | KRAS mutations status | Molecular Subtype | Tumor Site | AJCC Stage | Differentiation |
| --- | --- | --- | --- | --- | --- |
| **SNU-410** | KRAS_G12D | Squamous |  |  |  |
| **PK45H** | KRAS_G12D | Squamous |  |  |  |
| **PL45** | KRAS_G12D | Squamous |  |  |  |
| **PaTu 8988T** | KRAS_G12V | Squamous |  |  |  |
| **PaTu 8988S** | KRAS_G12V | Progenitor |  |  |  |
| **HPAC** | KRAS_G12D | Progenitor |  |  |  |
| **HPAFII** | KRAS_G12D | Progenitor |  |  |  |
| **ASPC-1** | KRAS_G12D | Progenitor |  |  |  |
| **MDA-PATC102** | HRAS_G12D | Progenitor | Pancreatic body | IV | Poor |
| **MDA-PATC108** | KRAS_G12D | Progenitor | Pancreatic head | IIA | Poor |
| **MDA-PATC148** | KRAS_G12D | Squamous | Liver | IV | n/a |
| **MDA-PATC153** | WT | Squamous | Pancreatic body | IIB | Poor |

